# Selective proteasomal degradation from the Golgi apparatus membrane

**DOI:** 10.64898/2026.02.04.703791

**Authors:** Joao Diamantino, Sofia Murteira, Steffen Lawo, Farnusch Kaschani, Markus Kaiser, Doris Hellerschmied

## Abstract

To maintain cellular homeostasis, protein quality control (PQC) machineries target damaged proteins for degradation. However, the PQC machinery, operating at the major protein sorting and processing site of mammalian cells, the Golgi, has remained largely elusive. Here we used a chemical biology tool to induce misfolding and ubiquitination of Golgi-localized model substrates. Our tool recruits a Cullin-RING ubiquitin ligase complex by exposure of a degron, recognized by the substrate receptor KLHDC2. Using this tool, we found that ubiquitinated Golgi membrane proteins are targeted for proteasomal degradation, a process that is critical to avoid Golgi fragmentation. This process is facilitated by the p97-unfoldase, which assembles with its adaptors UFD1-NPL4 and FAF2 at the Golgi. We further identified the rhomboid pseudo-protease RHBDD2 as an important contributor to Golgi homeostasis as it binds misfolded, ubiquitinated Golgi membrane proteins and regulates Golgi morphology. Our findings establish a molecular framework for Golgi-PQC in mammalian cells.

- The accumulation of misfolded Golgi proteins causes Golgi fragmentation.
- Ubiquitinated Golgi membrane proteins are degraded by the proteasome.
- The p97 unfoldase operates at the Golgi with its adaptors UFD1-NPL4 and FAF2.
- The rhomboid pseudo-protease RHBDD2 contributes to Golgi protein quality control.

## INTRODUCTION

The Golgi apparatus (Golgi) is a critical organelle in eukaryotic cells. As part of the secretory pathway, the Golgi together with the endoplasmic reticulum (ER) ensures the faithful production, maturation, and sorting of nearly all cellular secretory and transmembrane proteins.^1–3^ In mammalian cells, the Golgi consists of stacked membrane-enclosed compartments that are laterally linked to form the Golgi ribbon.^4^ Disruption of this structural organization, Golgi fragmentation, impairs protein trafficking and Golgi-localized signalling complexes.^5–8^ Moreover, Golgi fragmentation is a sign of cellular stress and linked to disease onset and progression. ^9–11^ For example, as a hallmark of certain neurodegenerative diseases, Golgi fragmentation leads to impaired protein sorting, driving disease-related proteotoxic stress.^12–15^

To maintain cellular homeostasis and avoid the formation of toxic protein aggregates, organelle-specific PQC machineries target unwanted, misfolded, or orphaned proteins for degradation.^16–18^ To this end, the ubiquitin-dependent clearance of proteins from the Golgi has been reported in budding yeast. In particular, the cytosolic ubiquitin ligase Rsp5 is recruited to target Golgi-localized substrates^19^ and thereby induces their vacuolar degradation, mediated by the ‘endosomal sorting complexes required for transport’ (ESCRT) machinery.^20,21^ Additionally, the multi-subunit ‘Defective in SREBP Cleavage’ (Dsc) complex, which is integral to the Golgi membrane, targets substrates for degradation.^22^ The Dsc2 subunit, a rhomboid pseudo-protease, uses its membrane thinning activity for substrate recognition^23^ and substrate ubiquitination is carried out by the ubiquitin ligase subunit, Tul1^24–26^. Select Dsc complex substrates are targeted for proteasomal degradation through the endosome and Golgi-associated degradation (EGAD) pathway.^23,27,28^ This pathway comprises the extraction of the ubiquitinated substrate from the Golgi membrane by the hexameric AAA+ ATPase Cdc48 (p97/VCP in mammalian cells), which is recruited by its co-factor and Dsc complex subunit, Ubx3 ^27^. Additionally, ubiquitin-dependent proteases can shed protein domains from the Golgi membrane to liberate them for subsequent vacuolar or proteasomal degradation.^29^

Despite its important role in Golgi PQC in yeast, the Dsc complex lacks Golgi-localized homologs in mammalian cells.^22,27^ However, the EGAD pathway is conceptually similar to the well-characterized ER-associated degradation (ERAD) pathway, conserved from yeast to mammals. Also at the ER, misfolded proteins are labeled with ubiquitin by transmembrane ubiquitin ligase complexes and extracted by the Cdc48/p97 unfoldase for subsequent proteasomal degradation.^30,31^ P97, like its yeast homolog Cdc48, requires co-factors and adaptors to regulate substrate targeting and its unfolding activity.^32^ Ubiquitinated substrates are recognized and engage p97 by the UFD1-NPL4 adaptor protein complex.^33^ Additionally, the accessory adaptor FAF2 facilitates the p97-dependent degradation of transmembrane and membrane-associated proteins.^34–40^ FAF2 inserts into endomembranes with a short hydrophobic hairpin structure and thus functions at the ER, on mitochondria, lipid droplets, and peroxisomes.^34–36,41,42^ Upon membrane extraction, chaperones and shuttle factors support the handover of substrates for degradation in the proteasome.^31,43^

Although the degradation of damaged Golgi proteins has also been reported for mammalian cells, our knowledge of the PQC machinery is still limited.^44–46^ The Golgi is in close contact with a major degradative organelle, the lysosome. Indeed, lysosomal targeting and degradation has been reported for proteins with aggregated, highly oligomerized Golgi lumenal domains and Golgi-resident glycosylation enzymes, upon loss of their membrane anchor.^47–51^ More broadly, the lysosomal turnover of Golgi membranes is orchestrated by receptor-mediated autophagy, Golgiphagy, and Golgi Membrane Degradation (GOMED), important pathways contributing to Golgi homeostasis.^52–55^ Of note, the stress-induced degradation of the Golgi-associated protein GM130 revealed a role for the proteasome in regulating Golgi structure and homeostasis.^56^ This finding suggested that also in mammalian cells, p97 and the ubiquitin-proteasome system may be critical to the disposal of misfolded Golgi proteins.^56^

Here we combined a chemical biology approach, enabling induced protein misfolding at the Golgi in mammalian cells, with CRISPR/Cas9 knock-out screening and proteomics. We thereby show that the accumulation of misfolded Golgi proteins induces Golgi fragmentation, highlighting the importance of protein quality control for Golgi homeostasis. Clearance of Golgi proteins is driven by degron-induced ubiquitination and degradation in the proteasome. Mechanistically, we furthermore define p97, its adaptor proteins UFD1-NPL4 and FAF2, and the rhomboid pseudo-protease RHBDD2 as molecular machinery managing misfolded, ubiquitinated Golgi membrane proteins.

## RESULTS

### Accumulating misfolded Golgi proteins leads to Golgi fragmentation

To gain mechanistic insight into post-ER protein quality control, we followed the fate of misfolded Golgi proteins. We generated model substrates displaying a HaloTag2 (HT2) domain on the cytosolic face of the Golgi (**Figure 1A**). This domain can be locally misfolded by addition of the hydrophobic tag HyT36.^57–60^ As Golgi-targeting sequences we used GRASP55, a Golgi matrix protein, associated with the membrane by myristoylation. Moreover, we chose the integral Golgi membrane protein TMEM115 with six predicted transmembrane helices. The two model substrates were expressed in a doxycycline-dependent manner in stable HEK293 Flp-In T-REx cells (referred to as HEK293 throughout the text). In imaging experiments, we first confirmed that the model substrates localize to the Golgi, as outlined by immunofluorescence staining of the Golgi marker GM130 (**Figure 1B** and **S1A**). We then quantified the levels of TMEM115- and GRASP55-HA-EGFP-HT2 via EGFP fluorescence by flow cytometry after cycloheximide treatment for blocking protein re-synthesis (**Figure S1B** and **S1C**). Using this assay, we found that HyT36-treatment for 5 h results in a ∼40% and ∼50% decrease of TMEM115- and GRASP55-HA-EGFP-HT2 levels, respectively (**Figure 1C**). To determine the degradation pathway involved, we complemented these measurements with a co-treatment with the proteasome inhibitor MG-132 or the lysosome inhibitor Bafilomycin A1 (BafA1) (**Figure 1C**). Inhibition of the proteasome with MG-132 counteracted the HyT36-induced degradation, while inhibition of lysosomal degradation by BafA1 did not (**Figure 1C**). With the established model system, we assessed the phenotypic effects of inhibiting the degradation of misfolded Golgi proteins, focusing on changes in Golgi morphology as an indicator of Golgi stress. Based on immunofluorescence staining of the Golgi marker GM130, we quantified the number of Golgi elements to assess Golgi fragmentation and we determined the Golgi compactness index to additionally monitor changes in Golgi shape.^61^ Treating GRASP55-HA-EGFP-HT2-expressing HEK293 cells with the proteasome inhibitor MG-132 led to a significant increase in Golgi elements and a corresponding decrease in Golgi compactness, reflecting Golgi fragmentation. This effect was further exacerbated when misfolding of the Golgi model substrate was induced by HyT36-treatment (**Figure 1D** and **1E**). MG-132 and/or HyT36 treatment of TMEM115-HA-EGFP-HT2-expressing HEK293 cells induced a similar pattern (**Figure 1F** and **1G**). Comparing the Golgi compactness index of GRASP55-HA-EGFP-HT2- and TMEM115-HA-EGFP-HT2-expressing cells suggested that already the overexpression of the transmembrane substrate induces moderate Golgi fragmentation. Collectively, these data demonstrate that the Golgi model substrates GRASP55-HA-EGFP-HT2 and TMEM115-HA-EGFP-HT2 undergo proteasomal degradation and, thus, that the clearance of misfolded proteins from the Golgi apparatus is required to maintain Golgi homeostasis.

**Figure 1.**
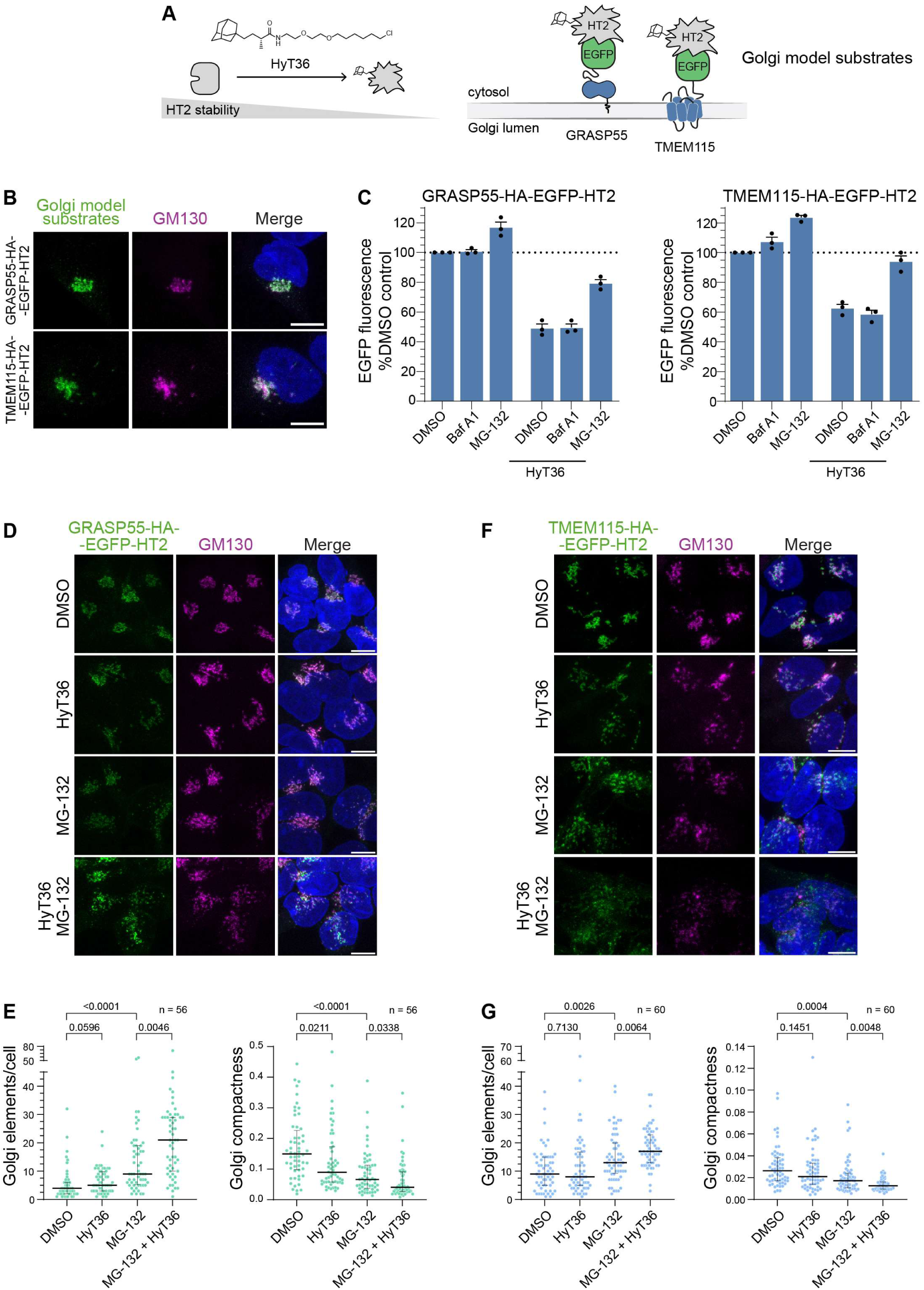
Protein misfolding at the Golgi membrane. (**A**) HaloTag2 (HT2) is misfolded upon HyT36 treatment. HT2-based model substrates are anchored to the Golgi membrane facing the cytosol. (**B**) Representative images of HEK293 cells expressing GRASP55-HA-EGFP-HT2 or TMEM115-HA-EGFP-HT2. Immunostaining of GM130 shows the Golgi. Scale bar, 10 µm. (**C**) Levels of GRASP55-HA-EGFP-HT2 or TMEM115-HA-EGFP-HT2 were analyzed by flow cytometry. Indicated treatments were applied for 5 h following a 1 h pre-treatment with cycloheximide. Data represent three independent experiments. Graphs show mean + S.D.(**D**) HEK293 cells expressing GRASP55-HA-EGFP-HT2 following the indicated treatments for 5 h were immunostained for GM130 to assess Golgi morphology. Images show representative maximum intensity projections of confocal image stacks. Scale bars, 10 µm. (**E**) Quantification of Golgi elements/cell and Golgi compactness based on the GM130 signal shown in (D). Representative results of two independent replicates. Error bars show median with interquartile range (IQR). P-value calculated with Kruskal-Wallis test (**F-G**) Same analysis as in (E-D) performed with HEK293 cells expressing TMEM115-HA-EGFP-HT2. Representative results of three independent replicates.

### Ubiquitination at the Golgi apparatus

We reasoned that TMEM115- and GRASP55-HA-EGFP-HT2 need to be ubiquitinated for degradation in the proteasome. To assess the ubiquitination state of the model substrates, we performed pull-down studies using Tandem Ubiquitin Binding Entities (TUBEs). With a TUBE construct comprising four tandem UBA^UBQLN1^ domains^62^, we enriched ubiquitinated proteins from HEK293 cell lysates and analyzed the elution fractions for the presence of TMEM115-and GRASP55-HA-EGFP-HT2 (**Figure 2A**). The TUBE pull-down showed that proteasome inhibition by MG-132 leads to a stabilization of the ubiquitinated TMEM115- and GRASP55-HA-EGFP-HT2 species, reflecting the natural turnover of the two model substrates by the ubiquitin-proteasome system. Under conditions of HyT36 treatment by itself we could not detect any accumulation of poly-ubiquitinated model substrates, as these were likely rapidly degraded. The strongest signal for the ubiquitinated species, was observed in samples treated with both, HyT36 and MG-132 (**Figure 2A**). We then used a ubiquitin-specific antibody (FK2)^63^ to determine the sub-cellular localization of ubiquitinated proteins (**Figure 2B** and **2C**). Upon proteasome inhibition with MG-132, we observed FK2-positive puncta throughout the cytoplasm and the nucleus of HEK293 cells expressing the Golgi model substrates. When we applied HyT36 to induce misfolding in combination with MG-132, we observed a significant increase of the FK2 signal coinciding with TMEM115- and GRASP55-HA-EGFP-HT2 in the Golgi area, as outlined by the Golgi marker GM130 (**Figure 2B** and **2C**). Collectively, these data demonstrate acute ubiquitination of the two model substrates at the Golgi upon HyT36-induced substrate misfolding.

**Figure 2.**
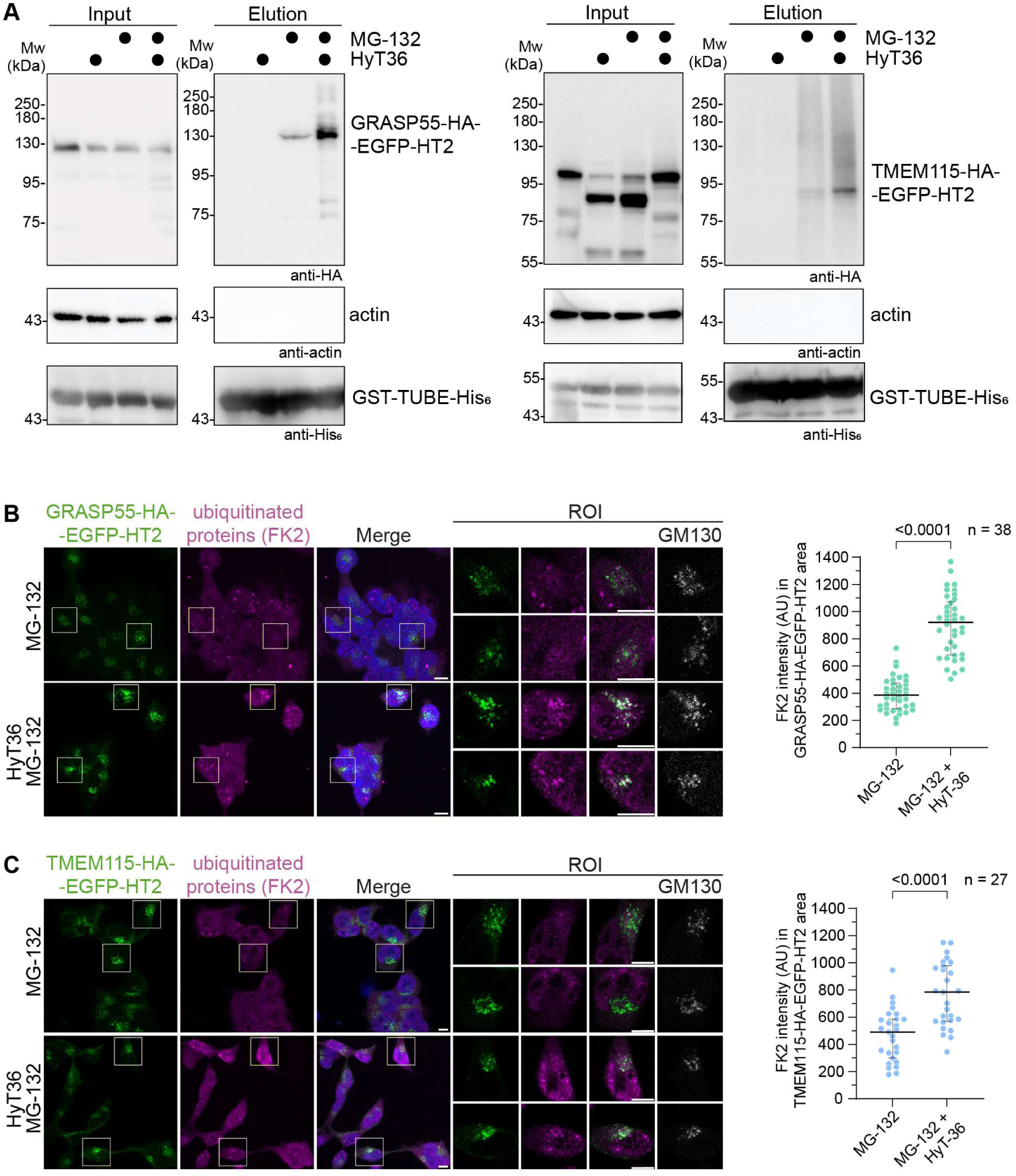
Ubiquitination of Golgi model substrates. (**A**) Tandem Ubiquitin Binding Entities (TUBE) pull-down of HEK293 cells expressing GRASP55-HA-EGFP-HT2 (left) or TMEM115-HA-EGFP-HT2 (right) following the indicated treatments for 5 h. Samples were analyzed by western blotting using the indicated antibodies. For TUBE pull-down of HEK293 cells expressing GRASP55-HA-EGFP-HT2, input and elution images were acquired using different exposure times. (**B**) *Left panel*: GRASP55-HA-EGFP-HT2 expressing HEK293 cells, which were treated with MG-132 or MG-132+HyT36 for 5 h were probed with FK2 antibody to visualize ubiquitinated proteins. Overview images show representative maximum projections of confocal image stacks. Confocal sections of regions of interest (ROIs) highlight the signal of GRASP55-HA-EGFP-HT2 and FK2 in the Golgi region as identified by GM130. Scale bar, 10 µm. *Right panel*: Quantification of FK2 signal in the GRASP55-HA-EGFP-HT2 positive area. Error bars show median with IQR. P-value calculated with Welch’s t-test. Representative results of two independent replicates. (**C**) Same analysis as in (B) for HEK293 cells expressing TMEM115-HA-EGFP-HT2.

### A C-degron pathway targets misfolded HT2 at the Golgi apparatus

To identify factors that facilitate the proteasomal degradation from the Golgi, we performed a pooled CRISPR/Cas9 knock-out (KO) screen using a ubiquitin-focused sgRNA library in HEK293 cells expressing GRASP55-HA-EGFP-HT2 (**Figure 3A**). Upon transduction with the sgRNA library, we treated the cells with doxycycline and the HT2-stabilizer HALTS1^64^ to accumulate GRASP55-HA-EGFP-HT2, and subsequently with cycloheximide to inhibit further protein synthesis. We then added HyT36 to induce the misfolding and proteasomal degradation of this GRASP55-HA-EGFP-HT2 pool. We hypothesized that genes whose knockdown enhances or inhibits proteasomal degradation from the Golgi would show a decrease or increase in EGFP fluorescence, respectively. We therefore isolated cells with high (top25) and low (bottom25) levels of GRASP55-HA-EGFP-HT2 by fluorescence activated cell sorting (FACS). These cells were subsequently analyzed by next generation sequencing and evaluated with the MAGeCK-RRA algorithm^65^ (**Figure 3B** and **Table S1**).

**Figure 3.**
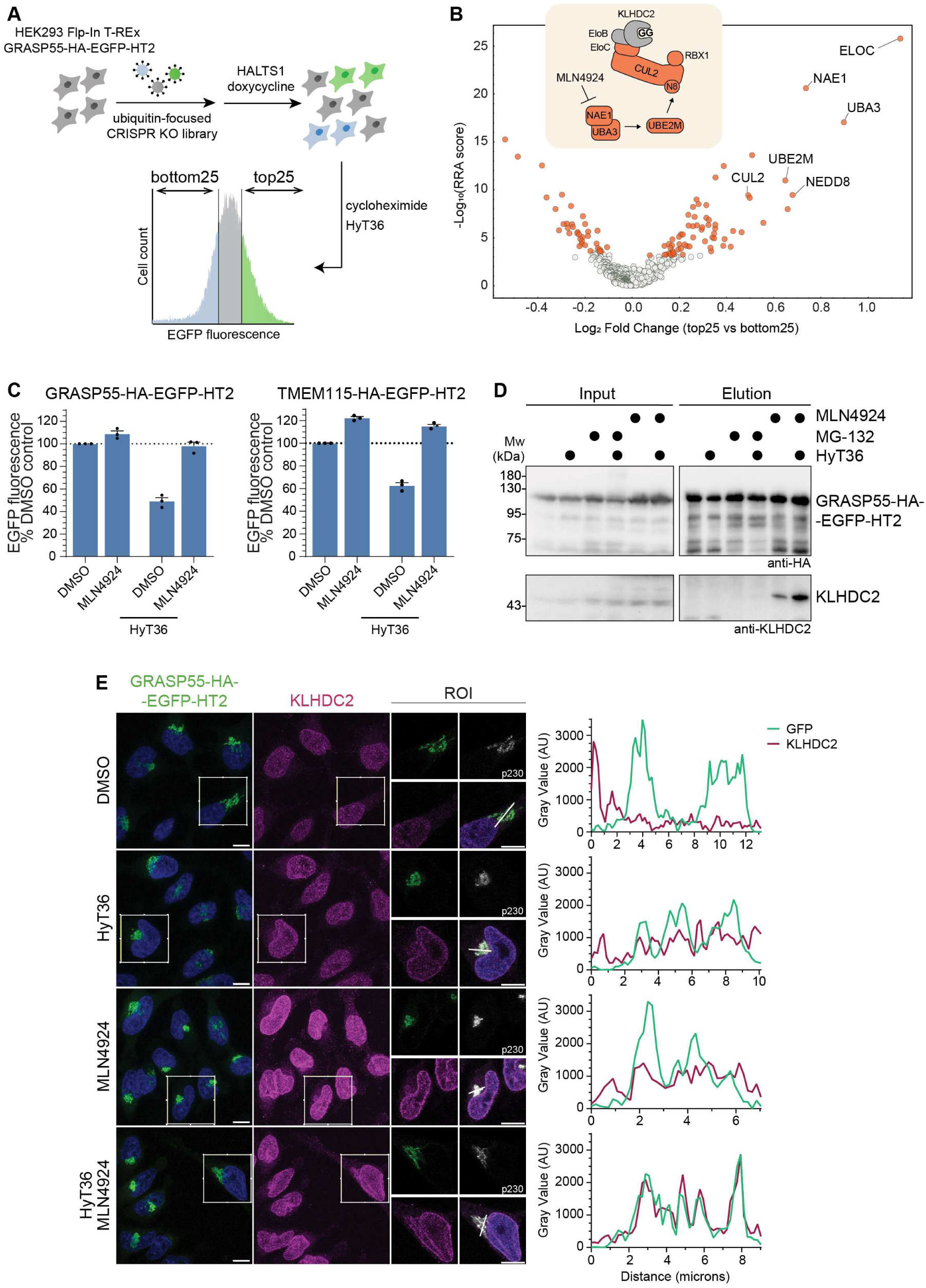
A Cullin-RING ubiquitin ligase targets Golgi model substrates. (**A**) Workflow of the CRISPR/Cas9 knock-out screen. (**B**) CRISPR/Cas9 screen results represented as volcano plot comparing genes targeted in the top25 and bottom25 cell populations. Significance value (RRA score) as calculated by the MAGeCK-RRA algorithm is plotted on the y-axis. Hits with an FDR ≤ 0.05 are highlighted in orange. NEDD8 (N8). (**C**) Levels of GRASP55-HA-EGFP-HT2 or TMEM115-HA-EGFP-HT2 were analyzed by flow cytometry. Indicated treatments, including the neddylation inhibitor MLN4924, were applied for 5 h following a 1 h pre-treatment with cycloheximide. Data represent three independent experiments. Graphs show mean + S.D. (**D**) GFP-trap pull-down of GRASP55-HA-EGFP-HT2 from HEK293 cells treated with the indicated compounds for 5 h. Input and elution fractions were analyzed by western blotting using the indicated antibodies. (**E**) *Left panel:* Overview images show representative maximum projections of confocal image stacks. Representative confocal sections of HeLa Flp-In T-REx cells expressing GRASP55-HA-EGFP-HT2. Following the indicated treatments for 5 h, cells were immunostained for KLHDC2 and the Golgi marker p230. *Right panel:* Quantification of the indicated gray value along the line shown in the region of interest (ROI). Scale bar, 10 µm. Arbitrary units (AU)

Six primary hits (ELOC, UBA3, NAE1, UBE2M, CUL2, NEDD8) from the screen suggested that GRASP55-HA-EGFP-HT2 is targeted by a multi-subunit Cullin-RING ligase (CRL) complex (**Figure 3B**). CRLs are activated by modification with the small protein modifier NEDD8 by an enzymatic cascade comprising NAE1/UBA3 and UBE2M.^66^ Accordingly, their activity can be inhibited using the neddylation (NAE1) inhibitor MLN4924.^67^ Hence, MLN4924 nearly abolished HyT36-induced degradation of TMEM115- and GRASP55-HA-EGFP-HT2 (**Figure 3C**). The hit with the highest log_2_-fold enrichment, ELOC, encodes the Elongin C (EloC) protein, which is part of CRL2 complexes (**Figure 3B**). Specifically, EloC and EloB (not targeted by the focused CRISPR library) form an adaptor protein complex, which bridges the Cul2 scaffold protein and BC-box substrate receptors.^68^ Among the approximately 40 different BC-box proteins, nine were shown to specifically bind C-terminal destabilizing motifs, known as C-degrons.^69,70^ Unlike the recognition of exposed hydrophobic residues in misfolded substrates, these Cul2-EloB/C-substrate receptor complexes identify short amino acid sequences at the C-termini of polypeptide chains, initiating their ubiquitination and subsequent proteasomal degradation.^69,70^ At the C-termini of TMEM115- and GRASP55-HA-EGFP-HT2, we identified a Gly-Gly (–GG) motif, which is recognized by the KLHDC2 substrate receptor.^69,70^ Of note, KLHDC2 was not targeted by the focused CRISPR library. Binding of a C-terminal –GG motif to KLHDC2 requires a total degron length of five amino acids in an extended conformation.^71,72^ The C-terminal glycines and the carboxyl group bind deep in the KLHDC2 Kelch repeat β-propeller domain, while the peptide backbone of the preceding three residues engages KLHDC2 residues lining the entry site of the binding pocket^71,72^ (**Figure S2A**). The C-terminal residues of HT2 are folded into an alpha helix (**Figure S2B**). However, HyT36 treatment induces unfolding of alpha helices^60^, offering a mechanism for HyT36-responsive exposure of the –GG degron. In pull-down experiments, we detected an interaction of KLHDC2 with GRASP55-HA-EGFP-HT2 in cells treated with MLN4924, likely reflecting that a pool of the model substrate is destabilized under steady-state conditions (**Figure 3D**). Consistent with our hypothesis, HyT36-induced misfolding of GRASP55-HA-EGFP-HT2 strongly increased its interaction with KLHDC2 (**Figure 3D**). To visualize the subcellular localization of this E3 ubiquitin ligase-substrate interaction, we performed immunofluorescence staining. Under control conditions in HeLa cells, KLHDC2 was found enriched in the nucleus and specifically at the nuclear membrane (**Figure 3E**), consistent with previous reports in different cell lines.^73^ When we however treated cells with HyT36 and MLN4924 to stabilize the interaction between TMEM115- or GRASP55-HA-EGFP-HT2 and KLHDC2, we found KLHDC2 localized to the Golgi, as outlined by the marker proteins p230 or GRASP65 (**Figure 3E** and **S2C**), suggesting its recruitment to the Golgi upon HT2 misfolding. By analyzing line profiles, we identified a distinct overlap between signals of the Golgi model substrates and KLHDC2 upon co-treatment of HyT36 and MLN4924 (**Figure 3E** and **S2C**). These data support the assembly of the CRL2^KLHDC2^ complex at the Golgi to induce the ubiquitination and degradation of EGFP-HT2 fusion proteins.

### Differential response to protein misfolding at cellular membranes

Our data showed that the CRL2^KLHDC2^ complex specifically responds to the HT2-hydrophobic tagging tool. Accordingly, HyT36 treatment produces two signals – a misfolded protein domain ^57,58,60^ and exposure of the C-terminal –GG degron. To discriminate the significance of both signals, we explored differences between membrane-associated and soluble cytosolic HT2 in mammalian cells. In addition to the Golgi-localized substrates, we generated HEK293 cell lines expressing soluble cytosolic HA-EGFP-HT2 and plasma-membrane localized FZD1-HA-EGFP-HT2 (**Figure S3A** and **S3B**). To assess the contribution of CRLs to the degradation of these EGFP-HT2 fusion proteins, we compared their degradation profile using MLN4924. We observed that, as for the Golgi-localized EGFP-HT2 proteins (**Figure 3C**), the plasma membrane-anchored FZD1-HA-EGFP-HT2 required the activity of a CRL for efficient HyT36-induced degradation (**Figure 4A**). In contrast, HyT36-induced degradation of soluble cytosolic HA-EGFP-HT2 was only mildly impaired by MLN4924 (**Figure 4A**). To further characterize this difference, we generated HEK293 cell lines expressing EGFP-HT2 fusion proteins terminating in a double aspartate sequence (–DD), preventing an interaction with KLHDC2 and promoting protein stability.^69^ Notably, a –DD sequence at the C-termini of TMEM115- and GRASP55-HA-EGFP-HT2 strongly impaired their HyT36-induced degradation. Indeed, 5 h after HyT36 treatment, protein levels of the –DD constructs were still comparable to the DMSO control (**Figure 4B**). The cytosolic HA-EGFP-HT2 and HA-EGFP-HT2^DD^ were however efficiently degraded, with a decrease of ∼55% and ∼35% after 5 h of HyT36-treatment, respectively (**Figure 4B**). Together with the observation of a persistent degradation of HA-EGFP-HT2 in presence of MLN4924 (**Figure 4A**), these data pinpoint a CRL-independent degradation of HA-EGFP-HT2. While misfolding is an efficient signal for inducing HT2 degradation in the cytosol, the degradation of misfolded Golgi-localized HT2 strongly depends on the exposure of the C-terminal degron sequence. To explore conditions that would favor degradation of misfolded Golgi proteins, we first monitored their levels upon extended treatment with HyT36. After 8 h, we observed a ∼5% and ∼10% decrease of TMEM115- and GRASP55-HA-EGFP-HT2^DD^, respectively (**Figure 4C**), suggesting that alternative, GG-degron-independent degradation mechanisms are activated at a later stage. Previous work demonstrated that misfolded HT2 interacts with Hsp70/90-type chaperones.^57,60,74^ We therefore explored whether inhibition of Hsp90 would impact protein degradation using the small molecule Hsp90 inhibitor 17-DMAG^75^. Co-treatment with 17-DMAG led to an increase in HyT36-induced degradation specifically of the –DD constructs, demonstrating that these misfolded Golgi proteins are sensitized to degradation in a challenged proteostasis environment (**Figure 4C**). Notably, also this degradation is proteasome-dependent (**Figure S3C**). Collectively, these data point to distinct mechanisms of substrate recognition on cellular membranes compared to the cytosol and highlight the importance of degrons in driving substrate ubiquitination on cellular membranes (**Figure 4D**).

**Figure 4.**
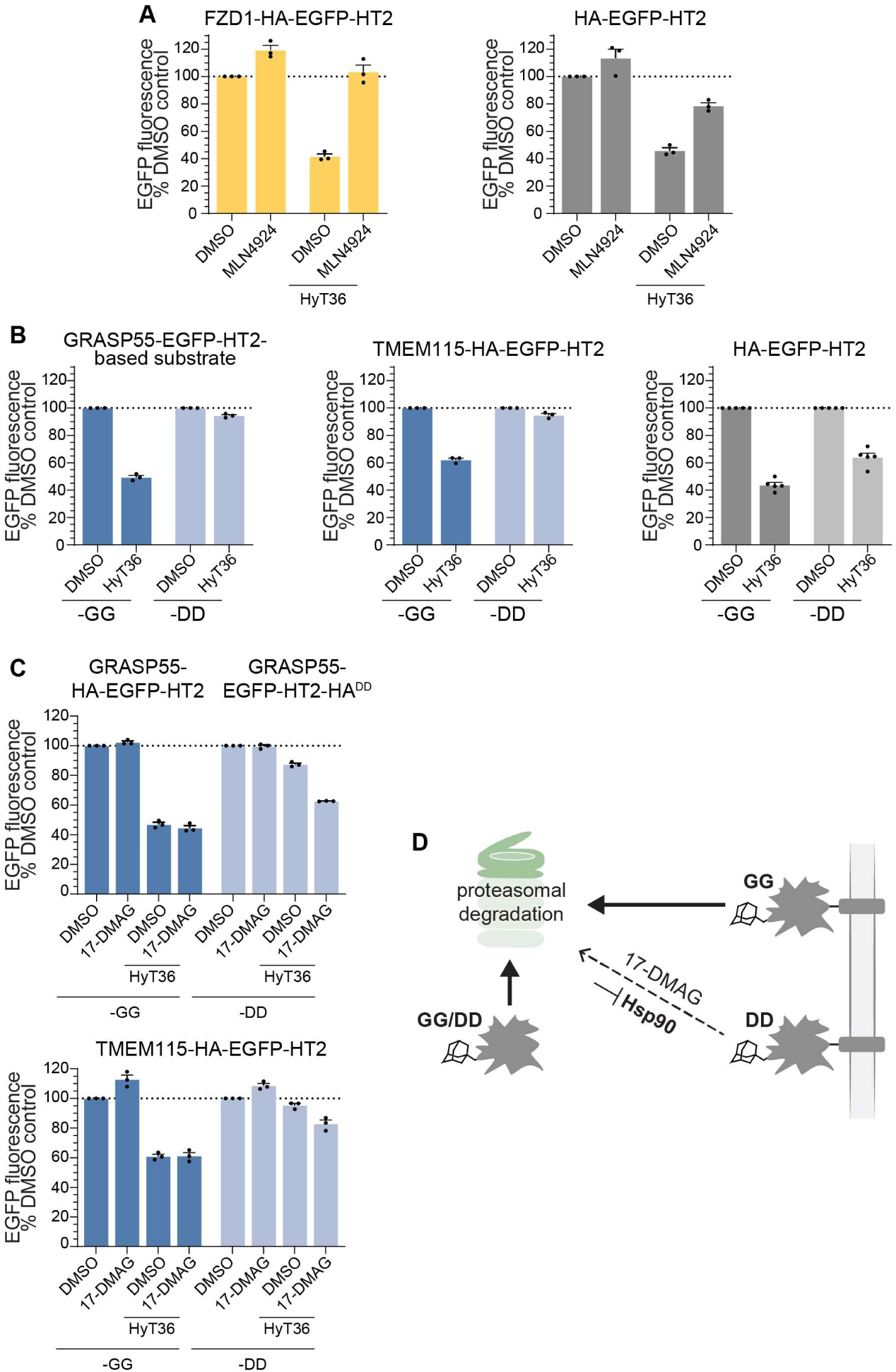
Comparative analysis of degradation from the Golgi membrane. (**A**) Levels of plasma membrane-localized FZD1-HA-EGFP-HT2 and cytosolic soluble HA-EGFP-HT2 were analyzed by flow cytometry. Indicated treatments, including the neddylation inhibitor MLN4924, were applied for 5 h following a 1 h pre-treatment with cycloheximide. (**B**) Comparison of the levels of the indicated EGFP-HT2 fusion proteins terminating in a –GG or –DD motif as determined by flow cytometry after DMSO or HyT36 treatment for 5 h following a 1 h cycloheximide pre-treatment. (**C**) Effect of the Hsp90 inhibitor 17-DMAG on the levels of the indicated Golgi proteins terminating in –GG or –DD, as determined by flow cytometry. Indicated treatments were applied for 8 h following a 1 h cycloheximide pre-treatment. (**D**) Summary of EGFP-HT2 degradation profiles. Misfolding-induced exposure of a GG-degron in membrane-anchored substrates leads to efficient degradation, while a C-terminal –DD sequence nearly abolishes degradation. Hsp90 inhibition by 17-DMAG sensitizes the membrane-anchored substrates terminating in –DD to degradation. In the cytosol, both substrate types i.e. terminating in –GG or –DD are efficiently degraded. For all graphs in this figure, data show at least three independent experiments. Graphs show mean + S.D.

### P97 inhibition enriches Golgi quality control complexes

We hypothesized that downstream of substrate recognition and ubiquitination, the AAA+ ATPase p97 contributes to the degradation of Golgi proteins. To explore the involvement of p97 in protein degradation, we used the small molecule p97 inhibitor CB-5083^76^. For GRASP55-HA-EGFP-HT2 and HA-EGFP-HT2, CB-5083 co-treatment led to a modest reduction in HyT36-induced degradation (**Figure 5A**), suggesting that p97 contributes moderately to the degradation of EGFP-HT2 constructs, possibly by supporting EGFP unfolding prior to its proteasomal degradation, as previously reported.^77,78^ Notably, for the integral Golgi membrane protein TMEM115-HA-EGFP-HT2, we found that its turnover and the HyT36-induced degradation strongly depend on p97 (**Figure 5A**). Under conditions of CB-5083 and HyT36 co-treatment, TMEM115-HA-EGFP-HT2 was stabilized at the Golgi as illustrated by the marker GRASP65 (**Figure S4A**).

**Figure 5.**
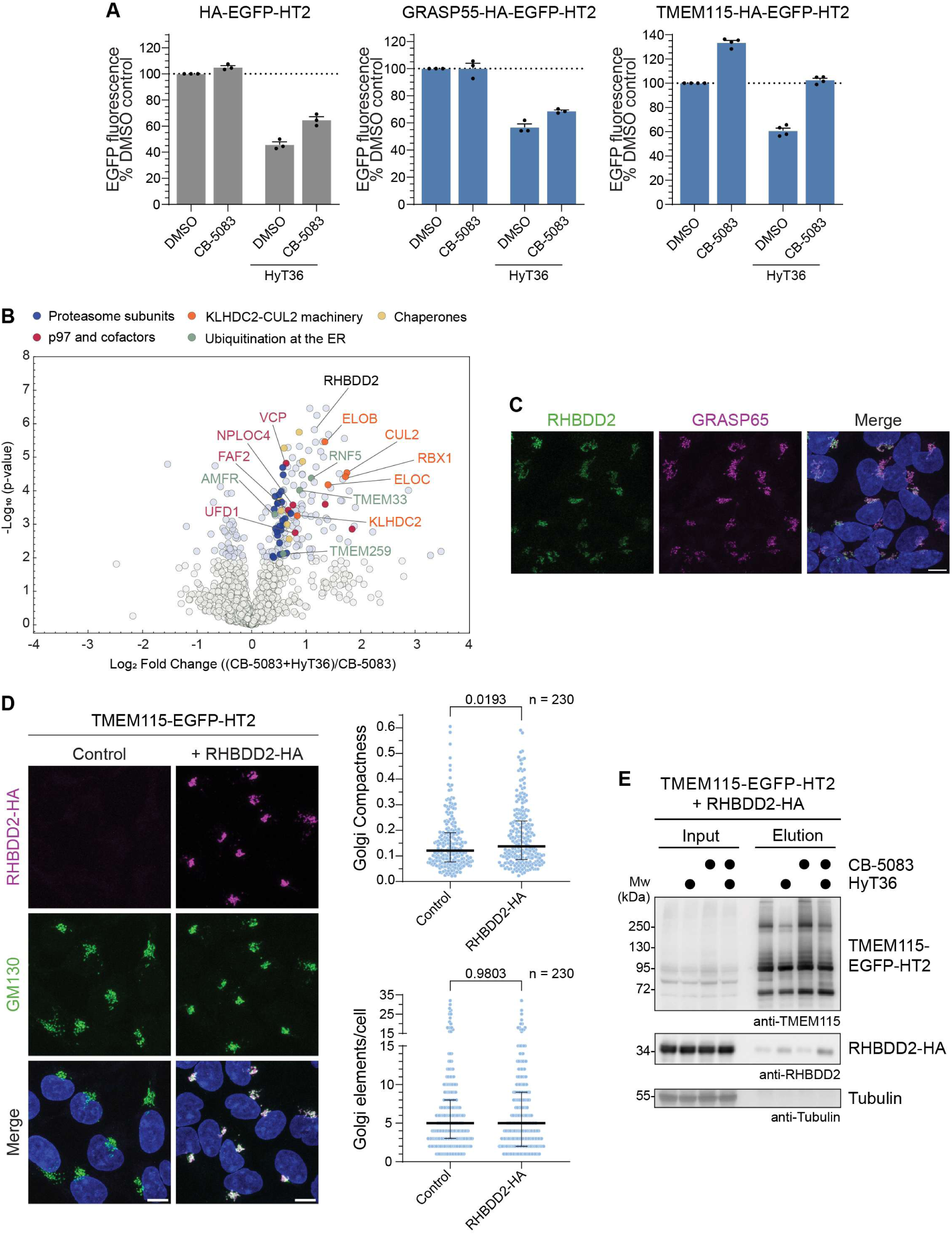
Inhibition of p97 stabilizes PQC machinery-substrate interactions. (**A**) Levels of the indicated EGFP-HT2 fusion proteins, as determined by flow cytometry, upon treatment with the p97 inhibitor CB-5083 and HyT36 for 5 h following a 1 h pre-treatment with cycloheximide. Data represent three independent experiments. Graphs show mean + S.D. (**B**) Volcano plot of proteins enriched by GFP-trap pulldown of TMEM115-HA-EGFP-HT2 from HEK293 cells comparing CB-5083 and CB-5083+HyT36 treatment conditions. Significantly altered proteins (p-value < 0.01) are shown in light blue. (**C**) Representative maximum intensity projections of confocal stacks of HEK293 cells immunostained for RHBDD2 and the Golgi marker GRASP65, showing the localization of the endogenous proteins. Scale bar, 10 µm. (**D**) *Left panel*: Representative maximum intensity projections of confocal stacks showing HEK293 cells with stable integration of TMEM115-EGFP-HT2 without its doxycycline induction. Additional stable integration of constitutively expressed RHBDD2-HA, detected by an HA-antibody, induces compaction of the Golgi, as visualized by immunostaining of GM130. Scale bar, 10 µm. *Right panel*: Quantification of Golgi elements and Golgi compactness in HEK293 cell lines shown on the left. Error bars show median with IQR. P-value calculated with Mann-Whitney U test. Representative results of two independent replicates. (**E**) GFP-trap pull-down from HEK293 cells expressing TMEM115-EGFP-HT2 and RHBDD2-HA treated with the indicated compounds for 3 h following a 3 h cycloheximide pre-treatment. Input and elution fractions were analyzed by western blotting using the indicated antibodies. Representative results of two independent replicates.

P97 is known to function with diverse adaptor proteins and integral membrane machinery in organelle-specific protein quality control pathways. We therefore hypothesized that protein quality control of TMEM115-HA-EGFP-HT2 at the Golgi requires additional components. Making use of our established, controllable chemical biology system to induce misfolding at the Golgi, we explored interactors of the TMEM115-HA-EGFP-HT2 substrate protein. We first pre-treated cells with cycloheximide for three hours, followed by an addition of either only the p97 inhibitor CB-5083 or CB-5083 together with HyT36 for three hours. We then performed a GFP-trap pull down and analyzed enriched TMEM115-HA-EGFP-HT2 binders via mass spectrometry (MS) (**Figure 5B** and **Table S2**). Importantly the two conditions yielded similar levels of TMEM115-HA-EGFP-HT2 (**Figure S4B**), thus allowing quantitative analyses of interactor proteins. Accordingly, we used label-free quantification (LFQ) to identify and quantify proteins specifically enriched under CB-5083+HyT36 treatment conditions. We observed an enrichment of different protein chaperones such as Hsp70 (*HSPA8*) and J-proteins (*DNAJA1*, *DNAJB2* – cytosolic, *DNAJB12* – ER-membrane-localized), consistent with the misfolded nature of the HT2 fusion protein upon conjugation to HyT36.^49,57,60,74^ We also found proteins with a well-established role in the ubiquitination of ERAD substrates (RNF5-TMEM33, Membralin, AMFR) among the enriched proteins, suggesting that despite cycloheximide treatment a fraction of TMEM115-HA-EGFP-HT2 is degraded from the ER in our pull-down set-up. Consistent with the results of the CRISPR/Cas9 KO screen and the *in cellulo* analysis (**Figure 3**), all subunits of the CRL2^KLHDC2^ ligase complex (CUL2, RBX1, EloC, EloB, KLHDC2), promoting ubiquitination at the Golgi were strongly enriched upon HyT36 treatment (**Figure 5B**). These data corroborate that the CRL2^KLHDC2^ complex targets TMEM115-HA-EGFP-HT2 at the Golgi.

In the proteomics data we further identified, RHBDD2 as an interactor of TMEM115-HA-EGFP-HT2 under conditions of CB-5083+HyT36 treatment. RHBDD2 belongs to the rhomboid pseudo-protease family. Rhomboid pseudo-proteases lack proteolytic activity, but have been implicated in the PQC of transmembrane proteins, including in collaboration with p97 ^79–81^. Dsc2, the rhomboid pseudo-protease component of the Golgi PQC machinery in yeast, mediates substrate recognition within the membrane.^23^ Other rhomboid pseudo-proteases in yeast and mammalian cells facilitate the removal of proteins from the ER membrane and subsequent proteasomal degradation.^82–84^ A structural model of RHBDD2 predicts the presence of a conserved rhomboid core consisting of six transmembrane helices and motifs previously described as important for membrane substrate removal (**Figure S4C**).^83^ Using an RHBDD2-specific antibody, we detected that the endogenous protein localized to the Golgi in HEK293 cells (**Figure 5C**). Given that the RHBDD2 antibody was of limited use in western blot experiments, we generated a HEK293 cell line co-expressing RHBDD2-HA and TMEM115-EGFP-HT2 to further explore their interaction. In these cells, the overexpressed RHBDD2-HA also localized to the Golgi (**Figure S4D**). In the imaging experiments, we further observed morphological changes of the Golgi upon over-expression of RHBDD2-HA. The Golgi, as visualized by the Golgi marker GM130 appeared smaller and more compact, suggesting that the overexpression of RHBDD2-HA resulted in a condensation of the Golgi structure (**Figure 5D**). Quantification of morphological features showed that the Golgi compactness index was significantly reduced by overexpression of RHBDD2-HA, while the number of Golgi elements was not affected, suggesting a particular influence on the shape of the Golgi (**Figure 5D**). From the stable cell line, we then isolated TMEM115-EGFP-HT2 by GFP-trap pull-down and observed that upon HyT36 treatment, RHBDD2 was enriched in the elution fraction (**Figure 5E**). This HyT36-induced interaction was stabilized by addition of the p97 inhibitor CB-5083, corroborating the results from the proteomics experiment. These data support that the Golgi-localized rhomboid pseudo-protease RHBDD2 contributes to Golgi PQC and that its activity impacts Golgi structure.

### The p97 adaptors UFD1-NPL4 and FAF2 cooperate with p97 in targeting Golgi membrane proteins

The proteomics data also reflected the p97-dependent proteasomal degradation of ubiquitinated TMEM115-HA-EGFP-HT2 (**Figure 5B**). Specifically, ubiquitin (RPS27A) and proteasome subunits were enriched upon HyT36 treatment (**Figure 5B**). The p97 inhibitor CB-5083 is known to induce trapping of substrate proteins in the p97 hexamer. Accordingly, we found that p97 as well as select p97 adaptors were enriched under CB-5083+HyT36 treatment conditions (**Figure 5B**). We focused our follow-up analyses on the UFD1-NPL4 adaptor protein complex and the accessory adaptor FAF2, given their previously reported activity in the degradation of ubiquitinated membrane proteins.^34,35,85^ Complementary to mass spectrometry, we found via western blot analysis that UFD1-NPL4 and FAF2 were enriched, particularly upon co-treatment of HyT36 with the p97 inhibitor CB-5083 and the proteasome inhibitor MG-132 (**Figure 6A**). The UFD1-NPL4 complex is required for p97-mediated processing of ubiquitinated substrates at diverse subcellular sites.^32,85^ FAF2 cooperates with UFD1-NPL4 in recruiting p97 to ubiquitinated membrane proteins and our data suggested that FAF2 also guides p97-dependent substrate processing at the Golgi membrane. To specifically visualize FAF2 in proximity of the Golgi-localized substrate protein TMEM115-HA-EGFP-HT2, we performed a proximity ligation assay (PLA). We detected endogenous FAF2 with a protein-specific antibody and TMEM115-HA-EGFP-HT2 with an anti-HA antibody. Following the PLA protocol, dot-like signals visualize proximity between FAF2 and the Golgi model substrate (**Figure 6B** and **S5**). In control conditions, in which we did not induce the expression of TMEM115-HA-EGFP-HT2, we detected a baseline background signal. Upon expression of TMEM115-HA-EGFP-HT2, the appearance of PLA dots indicated proximity to FAF2. Notably, following HyT36 treatment the PLA dots accumulated in proximity of the condensed TMEM115-HA-EGFP-HT2 structure, demonstrating translocation of FAF2 to the Golgi area (**Figure 6B**). Taken together, our data support that a p97-UFD1-NPL4-FAF2 complex operates at the Golgi to facilitate the proteasomal degradation of ubiquitinated Golgi membrane proteins.

**Figure 6.**
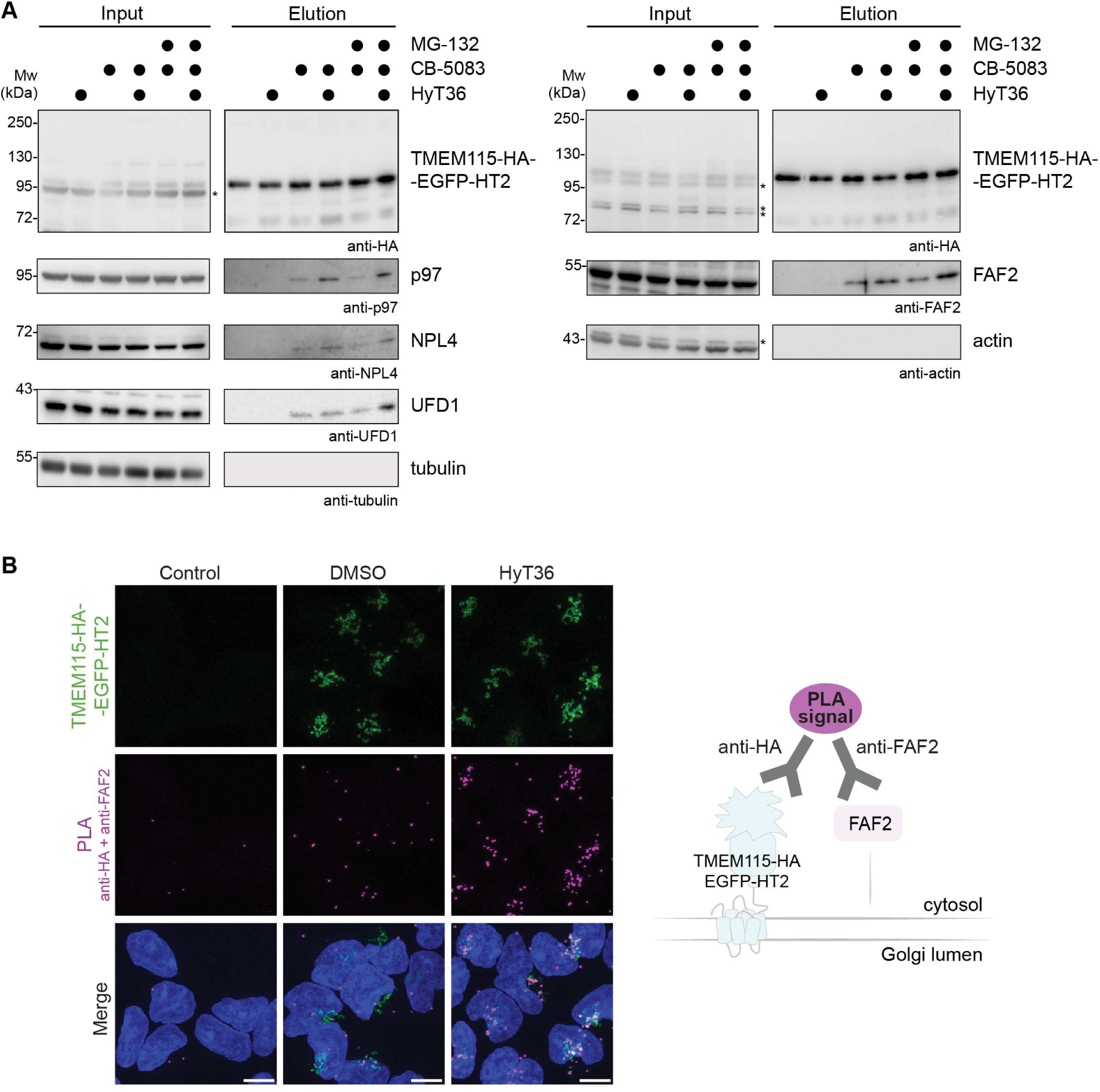
P97 adaptor proteins at the Golgi. (**A**) GFP-trap pull-down of TMEM115-HA-EGFP-HT2 from HEK293 cells treated with the indicated compounds for 3 h following a 3 h cycloheximide pre-treatment. Input and elution fractions were analyzed by western blotting using the indicated antibodies. Input and elution images were acquired using different exposure times. Representative results of two independent replicates. Asterisks mark signals from previous antibody incubations. (**B**) Proximity ligation assay (PLA) using FAF2- and HA-antibodies in three conditions. In control conditions, the expression of TMEM115-HA-EGFP-HT2 integrated into HEK293 cells was not induced. Doxycycline-induced HEK293 cells were treated with DMSO or HyT36 for 3 h following a 3 h cycloheximide pre-treatment.

## DISCUSSION

In this study, we uncovered molecular mechanisms underlying Golgi protein quality in mammalian cells, showing similarities to the EGAD pathway in yeast. Based on our results, we propose an emerging model for the selective proteasome-dependent degradation of proteins from the Golgi membrane, comprising their ubiquitination at the Golgi and p97-mediated extraction, supported by UFD1-NPL4 and FAF2 adaptors, as well as by the rhomboid pseudo-protease RHBDD2. This degradation is critical to avoid the accumulation of misfolded Golgi proteins, which compromises homeostasis by inducing Golgi fragmentation (**Figure 7**).

**Figure 7.**
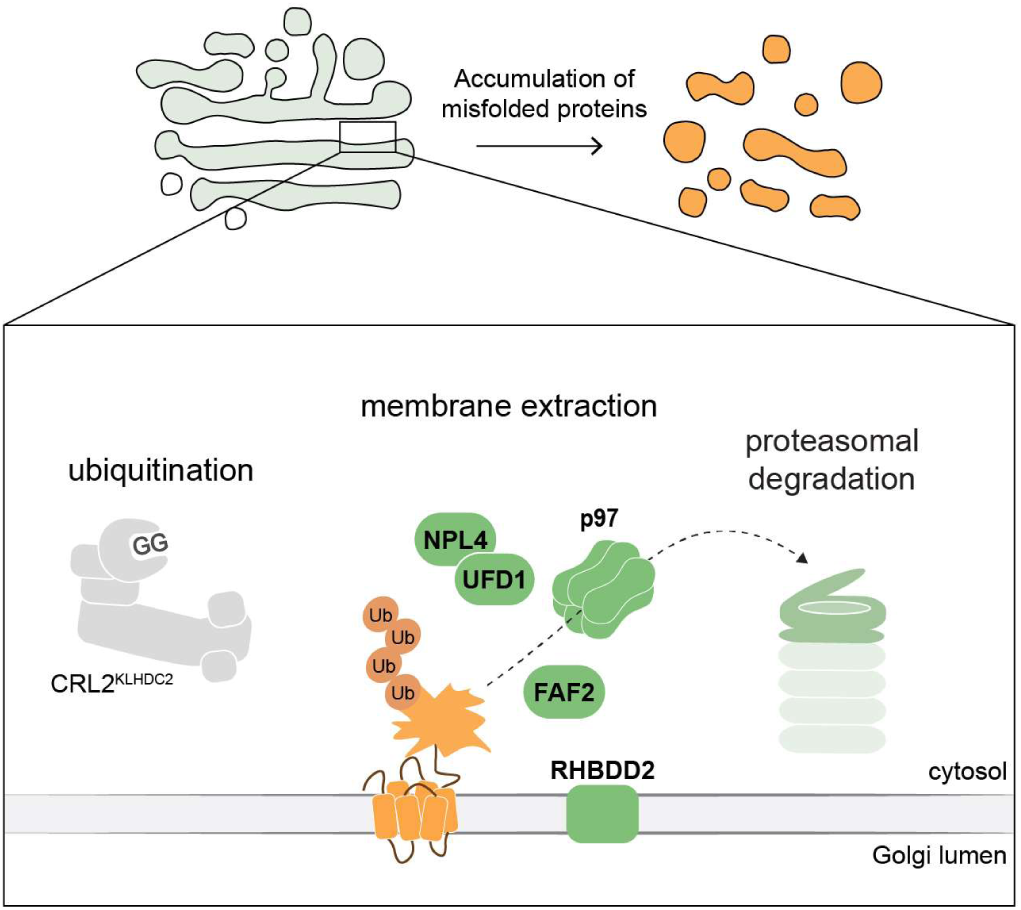
Model of the proteasomal degradation of Golgi membrane proteins in mammalian cells.

In our work, misfolding and ubiquitination of Golgi-localized substrates is achieved by the use HT2 fusion proteins and the chemical tool HyT36. We found that in the case of HT2, the folding state dictates the accessibility of a C-degron motif, highlighting a concept for distinguishing between folded and misfolded proteins beyond the exposure of hydrophobic segments. In a comparative analysis with a soluble HT2 fusion protein, we found that in the absence of a degron, HT2 misfolding at the Golgi or the plasma membrane is inefficient in targeting proteins for degradation. This finding is consistent with previous work in yeast comparing the processing of membrane-anchored and soluble misfolded protein domains by the PQC machinery.^86^ Misfolded domains localized in the cytosol or at the ER membrane are degraded via proteasome-dependent pathways, whereas those localized in post-ER compartments show minimal engagement with either proteasomal or vacuolar degradation mechanisms.^86^ Together these data highlight the need for alternative mechanisms of substrate recognition in protein quality control at post-ER membranes. Our data suggest that the exposure of the C-terminal degron of HT2 dominates substrate selection at cellular membranes. Interestingly, a role for the C-degron pathway in PQC of endogenous secretory and transmembrane proteins has previously been demonstrated.^87^ C-degrons are largely depleted from the eukaryotic proteome, except for transmembrane and secretory proteins.^87^ The accessibility of degrons can serve as an indicator of the topology of transmembrane proteins.^87^ Upon proper membrane insertion, the C-degrons would be hidden in the lumen of cellular organelles, contributing to the stabilization of transmembrane proteins. We speculate that in the context of endogenous Golgi proteins, misfolding and/or protease cleavage may expose degrons to engage PQC pathways. Moreover, we also identified a condition under which a degron-independent pathway was enabled. Inhibition of the Hsp90 chaperone machinery using 17-DMAG sensitizes misfolded Golgi-localized model substrates to proteasomal degradation, indicating that the visibility of misfolded substrates at the Golgi membrane is increased in a challenged proteostasis environment.

In addition to differences in substrate recognition, PQC pathways at cellular membranes face a challenge in accessing degradation pathways in lysosomes or the proteasome. In yeast, ubiquitinated Golgi membrane proteins can be targeted to the vacuole facilitated by the ESCRT machinery, or to the proteasome facilitated by Cdc48.^45,88^ In mammalian cells, we found that the degradation of the integral plasma membrane protein FZD1-HA-EGFP-HT2 depends partly on the proteasome and partly on lysosomal activity (**Figure S3B**), while ubiquitinated TMEM115-HA-EGFP-HT2 is degraded primarily by the proteasome. Our work specifically established the p97-dependent proteasomal degradation of transmembrane proteins from the Golgi (**Figure 7**). This aligns with a previously described role of p97 in the degradation of the Golgi-associated protein GM130.^56^ We further identified an interaction with the p97 adaptor protein complex UFD1-NPL4 that harbors a well-established role in targeting ubiquitinated substrates^32,85^, and the accessory adaptor FAF2. Recent work showed a role for FAF2 in targeting ubiquitinated proteins across a range of cellular membranes.^34–36,38^ The membrane-inserting hairpin of FAF2 confines p97 activity, while at the same time the C-terminal UBX domain and a preceding helix accelerate p97 activity.^89,90^ With these combined properties, the FAF2-induced boost in p97 activity may specifically provide the energy to support the extraction of transmembrane helices. Our data support that FAF2 contributes to the degradation of Golgi membrane proteins in mammalian cells, matching its proposed functional homolog Ubx3 in yeast^27^ and its role at other membrane-enclosed compartments.

In addition, membrane PQC is supported by rhomboid pseudo-proteases.^79–81^ Our data showed that upon induced misfolding and ubiquitination, TMEM115-EGFP-HT2 associates with RHBDD2, a poorly characterized rhomboid pseudo-protease. RHBDD2 shows the typical rhomboid protease fold (**Figure S4C**), but lacks the active site serine required for protease activity^91^. Rhomboid pseudo-proteases are thought to function by inducing local membrane lipid distortion and membrane thinning.^81^ In the EGAD complex, the rhomboid pseudo-protease subunit Dsc2 uses its membrane thinning activity to recognize orphaned proteins, which expose transmembrane segments typically hidden by binding partners, and induces their ubiquitination by Tul1.^23^ As part of ERAD complexes, members of the derlin family of rhomboid pseudo-proteases facilitate substrate removal from the ER membrane and coordinate subsequent proteasomal degradation.^82–84,92,93^ In a similar fashion, we propose a role for RHBDD2 in targeting substrates for proteasomal degradation, but – given its localization – from the Golgi membrane. The impact of RHBDD2 on Golgi structure, as observed by Golgi compaction upon its overexpression, further supports a role in Golgi homeostasis.

Our findings provide important insight into Golgi homeostasis mechanisms and motivate the search for additional PQC machinery such as ubiquitin E3 ligases, which may either be recruited from the cytosol or may be integral to the Golgi membrane and for endogenous substrates regulated by this EGAD-like pathway in mammalian cells.

## ACKNOWLEDGEMENTS

We thank Joel Brenner for the purification of the GST-TUBE-His_6_ protein and Emilia Chlosta, Rebecca Lehmann, Nina Marx, Maren Naudszus, and Rebecca Schulz, for support of initial cell biology and biochemistry experiments. We would like to thank Jenny Bormann and Svenja Heimann for excellent technical support of MS experiments. We acknowledge the Imaging Center Campus Essen (ICCE), Center of Biotechnology (ZMB), University of Duisburg-Essen, for providing the imaging equipment and support in microscope usage and image analysis. The Leica TCS SP8 HCS A microscope was funded by the Deutsche Forschungsgemeinschaft (DFG, German Research Foundation), Project-ID 219183055. The Leica TCS SP8X FALCON microscope was funded by the DFG, Project-ID 397277702. We thank Hemmo Meyer, Marius Lemberg, and Yevgeniy Serebrenik for critical reading of the manuscript. DH, FK, and MK were supported by the DFG (SFB1430, project 424228829). JD, SM, and DH were supported by the Sofja Kovaleveskaja Award by the Alexander von Humboldt Foundation endowed by the Federal Ministry of Education and Research.

## AUTHOR CONTRIBUTIONS

JD and SM performed cell biology and biochemistry experiments. JD performed imaging and image analysis. JD and SL performed the CRISPR screen. FK and MK performed MS experiments and analyzed the data together with SM. All authors contributed to data interpretation. JD, SM, and DH prepared the manuscript with input from all co-authors.

## EXPERIMENTAL PROCEDURES

### Reagents

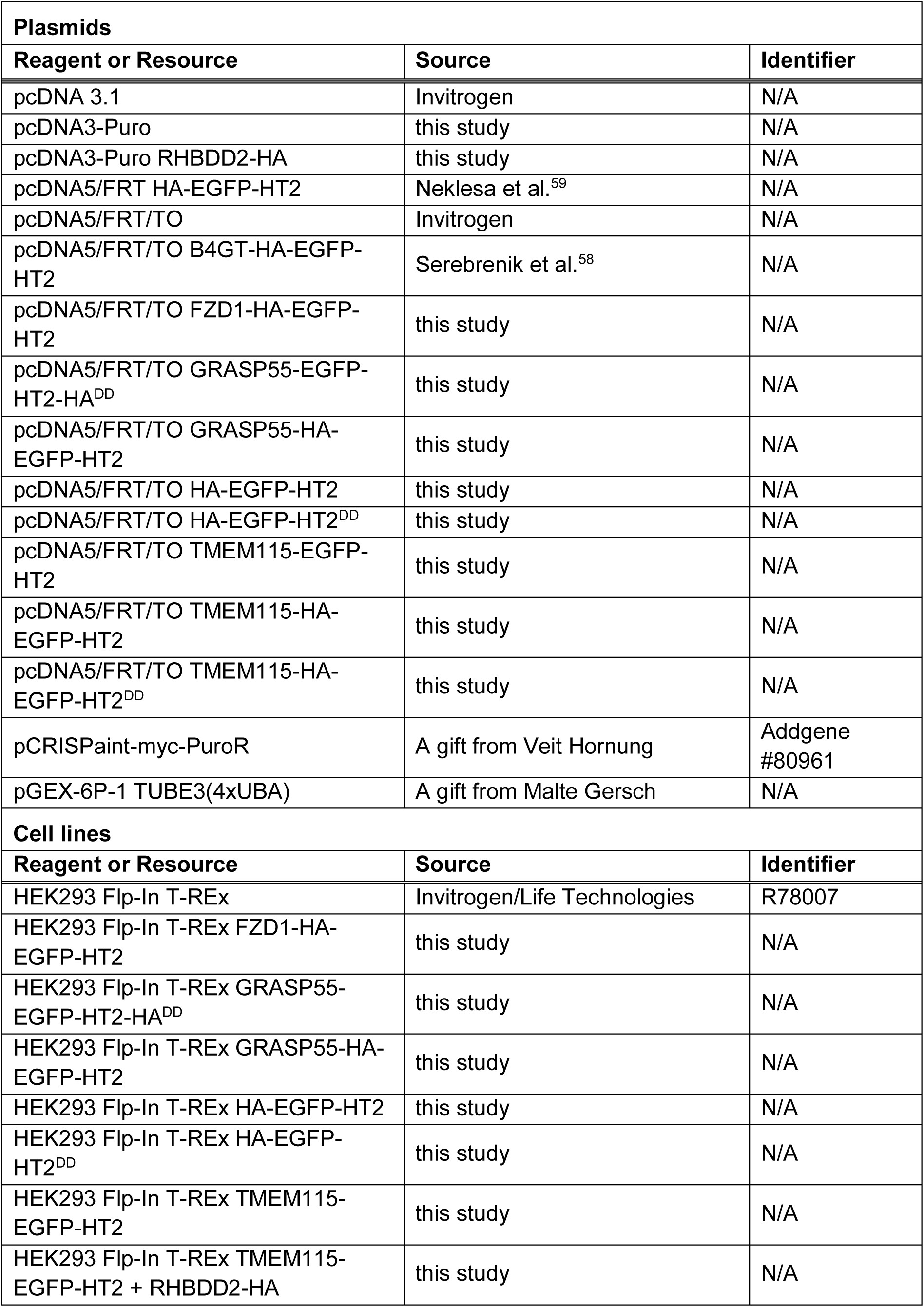
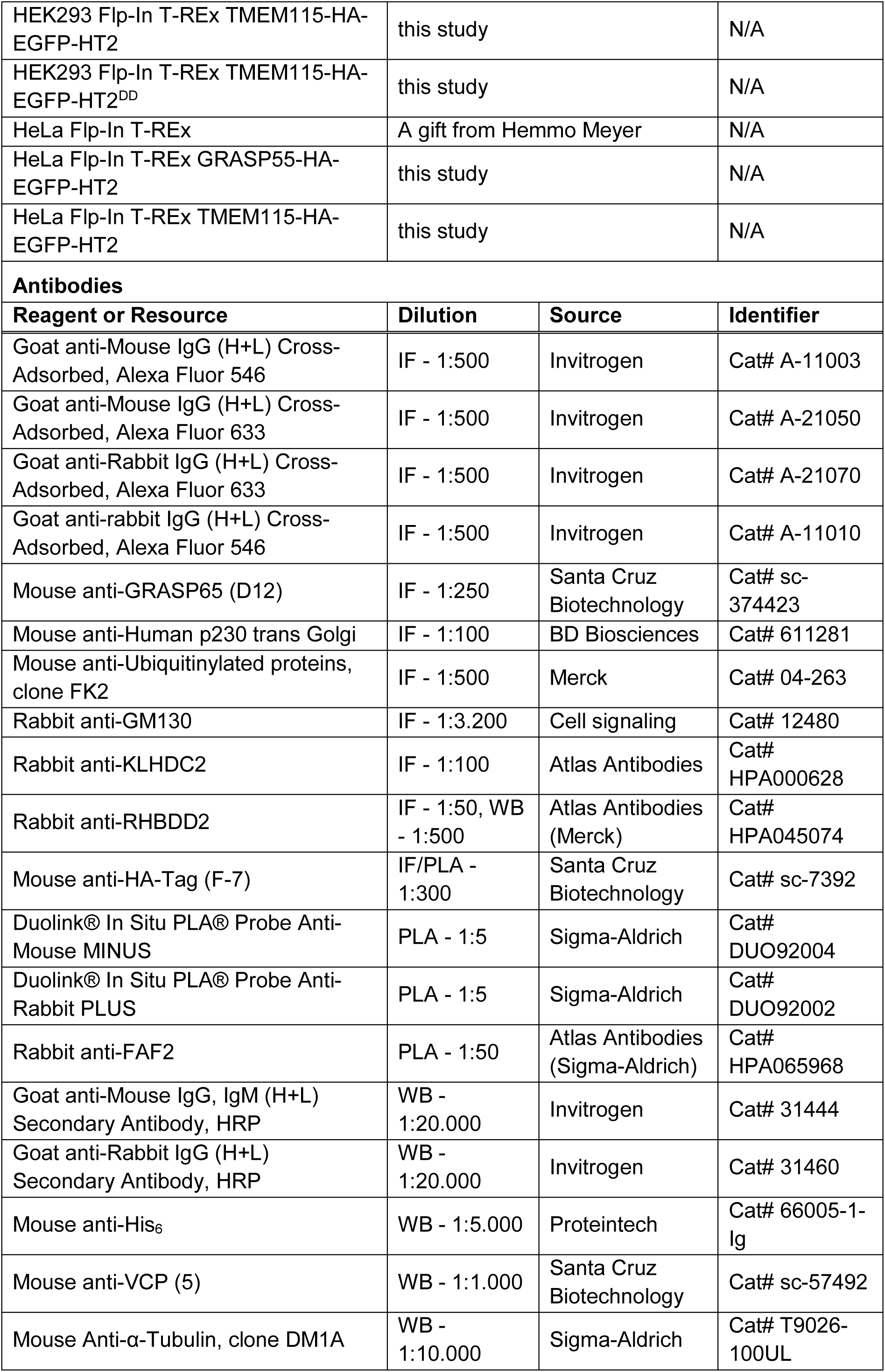
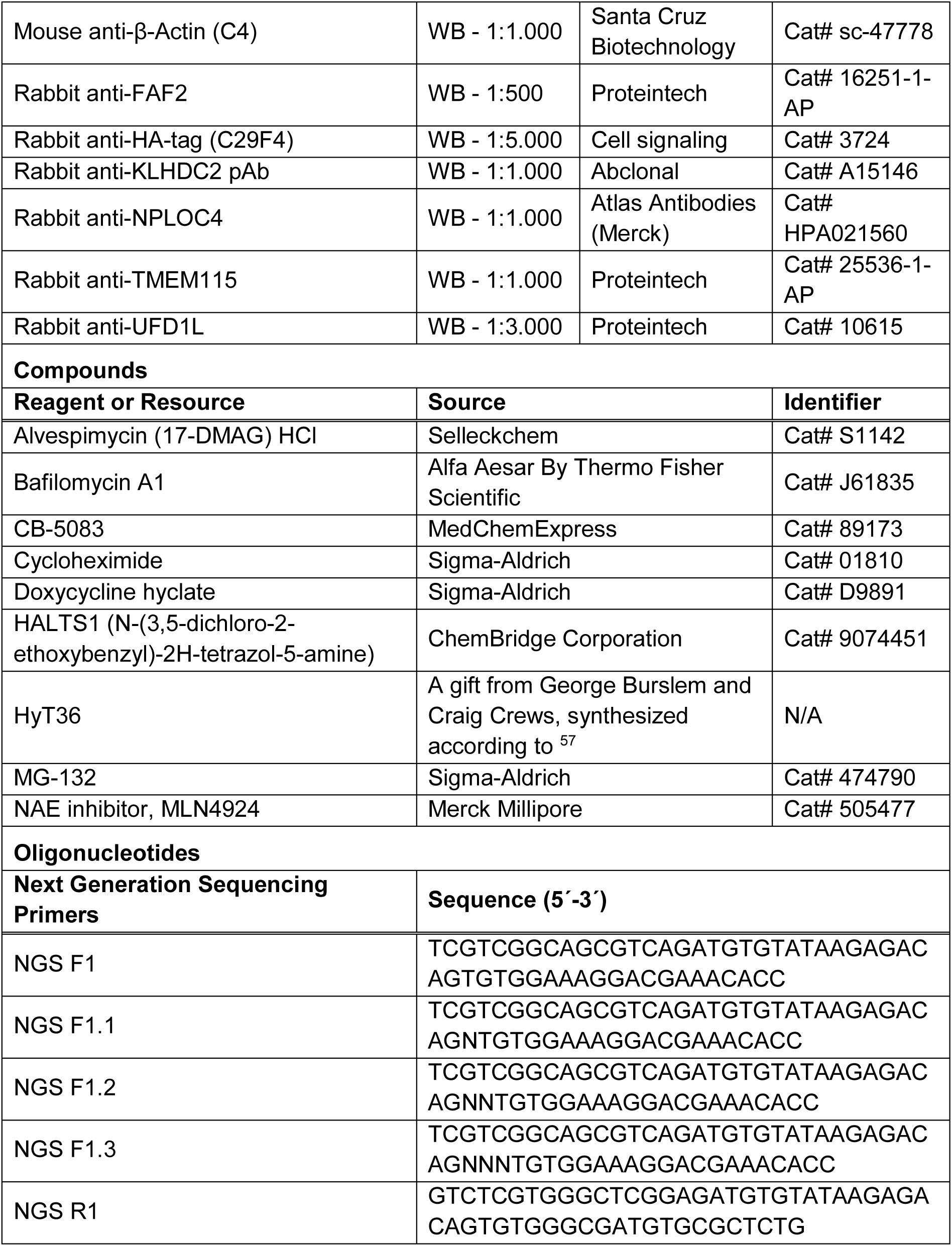

### Cloning

All DNA constructs were generated using the NEBuilder HiFi DNA Assembly Cloning Kit (NEB, E5520S). To generate pcDNA5/FRT/TO GRASP55-HA-EGFP-HT2, pcDNA5/FRT/TO TMEM115-HA-EGFP-HT2, or pcDNA5/FRT/TO FZD1-HA-EGFP-HT2, B4GT was removed by restriction digest (HindIII/AgeI) from pcDNA5/FRT/TO B4GT-HA-EGFP-HT2 described in Serebrenik et al. ^58^. GRASP55, TMEM115, and FZD1 were amplified from HEK293 cDNA with accordingly designed DNA primers and inserted into the digested vector. Cloning of pcDNA5/FRT/TO TMEM115-EGFP-HT2 involved the amplification of TMEM115 and EGFP-HT2 sequences from pcDNA5/FRT/TO TMEM115-HA-EGFP-HT2, excluding the HA-tag. The two fragments were inserted into pcDNA5/FRT/TO, digested with HindIII and NotI. All HT2 fusion constructs contained a –GG motif at the C-terminus. TMEM115-HA-EGFP-HT2^DD^ was amplified from pcDNA5/FRT/TO TMEM115-HA-EGFP-HT2 with a reverse primer exchanging the last two glycines of the HT2 sequence by two aspartates. TMEM115-HA-EGFP-HT2^DD^ was then inserted into pcDNA5/FRT/TO, digested with HindIII and NotI. pcDNA5/FRT/TO GRASP55-EGFP-HT2-HA-DD was cloned by introducing GRASP55 and EGFP-HT2-HA-DD into a pcDNA5/FRT/TO, digested with HindIII and NotI. GRASP55 and EGFP-HT2-HA-DD sequences were separately amplified from pcDNA5/FRT/TO GRASP55-HA-EGFP-HT2, the latter with a reverse primer including an HA-tag followed by two aspartate residues. pcDNA5/FRT/TO HA-EGFP-HT2 (containing a –GG motif at the C-terminus) and pcDNA5/FRT/TO HA-EGFP-HT2^DD^ were cloned by amplifying HA-EGFP-HT2 and HA-EGFP-HT2^DD^ sequences from pcDNA5/FRT HA-EGFP-HT2 previously described in Neklesa et al.^59^ and inserting them into a pcDNA5/FRT/TO, digested with HindIII and NotI. The reverse primer amplifying HA-EGFP-HT2^DD^ exchanged the last two glycines of the HT2 sequence by two aspartates. The pcDNA3-Puro vector was generated by replacing the Neomycin resistance gene in pcDNA 3.1 by restriction digest with SmaI and BstbI and following insertion of a puromycin resistance gene, amplified from pCRISPaint-myc-PuroR (a gift from Veit Hornung, Addgene plasmid # 80961). RHBDD2-HA was introduced into pcDNA3-Puro by digesting the vector with HindIII and BamHI, and inserting RHBDD2-HA amplified from cDNA.

### Cell culture

HeLa Flp-In T-REx cells (a gift from Hemmo Meyer) and HEK293 Flp-In T-REx cells (Invitrogen/Life Technologies) were grown in 5% CO_2_ at 37 °C in DMEM (Invitrogen/Life Technologies, 31966047) containing 1% Pen/Strep (Invitrogen/Life Technologies, 15140122) and 10% fetal bovine serum (Invitrogen/Life Technologies, 10500056), referred to as growth medium. Stable HEK293 Flp-In T-REx and HeLa Flp-In T-REx cell lines were generated using the Flp-In™ T-Rex™ System (Invitrogen/Life Technologies, K6500-01) in combination with the pcDNA5/FRT/TO vector according to the manufacturer’s instructions, and the Transporter 5 Transfection Reagent (Polysciences Europe GmbH, 26008-5). Stable HEK293 Flp-In T-REx and HeLa Flp-In-T-REx cell lines were selected and cultured in growth medium containing 15 µg/ml blasticidin (Invivogen, ANT-BL-1), and 100 µg/ml or 250 µg/ml Hygromycin B (Invivogen, ANT-HG-5), respectively. For generation of double-stable cells, HEK293 Flp-In T-REx cells with stable integration of TMEM115-EGFP-HT2 were transfected with RHBDD2-HA in pcDNA3-Puro and selected using 1.5 µg/ml puromycin (Invivogen, ANT-PR-1). All cell lines were regularly tested for mycoplasma infection using VenorsGeM OneStep mycoplasma detection kit (Minerva Biolabs, 11-8050).

### Cell treatments

To induce expression of the different EGFP-HT2 fusion proteins, stable HEK293 Flp-In T-REx and HeLa Flp-In T-REx cells were treated with 100 ng/µl doxycycline hyclate for 24 h, except for the double stable HEK293 Flp-In T-REx TMEM115-EGFP-HT2+RHBDD2-HA cells, where expression of TMEM115-EGFP-HT2 was induced with 1 ng/µl doxycycline for 24 h. Treatments with 10 μM HyT36, 10 μM MG-132, 10 μM CB-5083, 1 μM MLN4924, 500 nM 17-DMAG and 100 nM Bafilomycin A1 were performed for 5 h, unless stated otherwise. Cycloheximide was used at a final concentration of 20 μg/ml for the indicated treatment times.

### Flow cytometry

HEK293 Flp-In T-REx cells expressing the different EGFP-HT2 fusion proteins, following the indicated treatments, were trypsinised, resuspended in growth medium and collected by centrifugation at 800 g for 5 min at RT. Growth medium was removed and cells were resuspended in 1% FBS, 1 mM EDTA in PBS and transferred to a 96-well plate. GFP levels were quantified by flow cytometry at a MACS QUANT MG16 flow cytometer (Miltenyi Biotec). Data were analyzed in Kaluza (Beckmann Coulter) and plotted in Prism (Graphpad).

### CRISPR/Cas9 knock-out (KO) screen

The Bonifacino Lab Human ubiquitination-related proteins CRISPR KO library^94^, containing 11,108 sgRNAs targeting 660 genes together with Cas9 was a gift from Juan Bonifacino (Addgene #174592). Lentiviral particles were produced by transfecting the library DNA along with lentiviral packaging plasmids (psPAX2 and pMD2.G) into HEK293T cells using Lipofectamine 3000 (Invitrogen L3000015). The plasmids psPAX2 and pMD2.G were gifts from Didier Trono (Addgene plasmids #12260 and #12259). Lentiviral supernatant was collected 48 h post-transfection, filtered (0.45 µm), and stored at −80 °C. The entire screen was carried out at a greater than 300x coverage per library sgRNA. 1.07 × 10^8^ HEK293 Flp-In T-REx GRASP55-HA-EGFP-HT2 cells were transduced with the library lentivirus at a multiplicity of infection of 0.25 using spinfection. 2 × 10^6^ cells and lentiviral particles were seeded into 12-well plates in growth medium containing 8 μg/ml polybrene (Santa Cruz, sc-134220) and spun at 1,500 rpm for 2 h in a heated centrifuge (37 °C). Afterwards, cells were recovered in fresh growth medium and plated in 5-layer flasks. After 48 h of recovery, cells were selected for 3 days in growth medium containing 1 μg/ml puromycin (Invivogen, ANT-PR-1), followed by 2 days of cell expansion in growth medium. After cell expansion, two replicates, each containing 4.4 × 10^7^ cells, were seeded into T175 flasks. 24 h later, the medium was supplemented with 100 ng/μl doxycycline and 10 μM HALTS1 and cells were incubated for another 24 h. A minimum of 5 × 10^6^ cells were then treated with 20 μg/ml cycloheximide for 1 h, followed by HALTS1 washout and a treatment with 20 μg/ml cycloheximide and 10 μM HyT36 for 5 h. For the pre-sort cell population, 2.5 × 10^7^ cells per replicate were pelleted and stored at −80 °C. Pre-sort samples were used for screen quality control (representation of library sgRNAs and dropout of sgRNAs targeting essential genes, compared to their representation in the plasmid pool). The remaining cells were stained with Fixable Viability Dye eFluor™ 780 (Invitrogen, 65-0865-18) and fixed with 2% Formaldehyde. After fixation, cells were washed with PBS and resuspended in PBS, 1% FBS, 1 mM EDTA. Cells were then sorted on two cell sorters in parallel (BD FACS ARIA IllU and BD FACS Aria Fusion) into gates with high and low EGFP signal, corresponding to the top and bottom 25% of the distribution of the EGFP fluorescence (top25 and bottom25 gate). Cells were collected in PBS, 2% FBS, 1 mM EDTA at 4 °C using a 100 μm nozzle with sheath pressure set at 20 p.s.i. into tubes previously coated with 20% FBS. Collected cells were pelleted and stored at −80 °C. Sorted cells were de-crosslinked in Buffer ATL (Qiagen 19076) with 2 mg/ml Proteinase K (Qiagen 19133) for 1 h at 56 °C, followed by 1 h at 90 °C. Genomic DNA was extracted using QIAamp DNA Blood Mini (Qiagen 51185) or Maxi Kits (Qiagen 51194) according to the manufacturer’s instructions. Following gDNA extraction, sgRNAs were amplified by PCR (24 cycles with 2.5 µg gDNA per reaction) with NEBNext UltraII Q5 DNA polymerase (NEB M0554L) and a mix of forward primers (NGS F1 – F1.3) and a reverse primer (NGS R1). PCR amplicons were pooled, bead-purified, quantified, and subjected to a second round of PCR to introduce Illumina Nextera adaptors and indices. The final library was bead-purified, quantified, pooled and analyzed by paired-end sequencing (2×100 bp) on an Illumina NovaSeq platform with >5 × 10^6^ reads per sample. After demultiplexing, raw NGS libraries were quality-checked using FastQC version 0.11.8 (Babraham Institute, Cambridge, UK)^95^. Upstream sequences and sgRNA length were used to trim reads with cutadapt (version 4.5). MAGeCK (version 0.5.9.5)^65^. was used to quantify the number of reads per sgRNA. Raw sgRNA counts were median-normalized, and MAGeCK test was used to rank sgRNAs. The log2-fold change (LFC) at the gene level was calculated with the median of the LFCs at sgRNA level. For gene significance, an α-RRA score and false discovery rate (FDR) were calculated by MAGeCK-RRA and plotted as a double-sided volcano plots using Instant Clue software v.0.11.3^96^.

### Fluorescence microscopy

HEK293 Flp-In T-REx cells were plated on coverslips coated with Poly-D-Lysine (Gibco. A38904-01), while HeLa Flp-In-T-REx cells were plated on coverslips coated with Fibronectin (Sigma-Aldrich, F0895). After the indicated treatments, cells were fixed using 4% formaldehyde in PBS for 15 min. Cells were permeabilized and blocked using PBS, 10% FBS, 0.1% Triton X-100, for 45 min at RT, followed by primary antibody (1 h at RT) and secondary antibody (45 min at RT) incubations. Antibodies were diluted in PBS, 1% BSA, 0.01% Triton X-100. Finally, nuclei were stained with 5 μg/ml Hoechst (Invitrogen, H1399) in PBS for 20 min at RT. Coverslips were mounted onto microscopy slides using Vectashield mounting medium (Vector Laboratories, VEC-H-1000). Microscopy images were acquired on a confocal laser scanning microscope (Leica TCS SP8 HCS A or Leica SP8X Falcon) equipped with PMT and HyD detectors and an HC PL APO 63x/1.4 CS2 objective. Images were acquired as z-stacks spanning the entire cell volume with a z-step size of 0.5 µm.

### Image analysis

Imaging data was quantified using Fiji^97^. The Golgi compactness index was determined as described in Bard et al^61^. Briefly, maximum intensity projections of the GM130 signal were generated and background pixels were removed by using the Otsu threshold. A region of interest was drawn around a single Golgi and Golgi elements were identified (minimum area of 0.05 µm^2^). The area and perimeter of each Golgi element within the region was determined and the compactness was calculated with the formula 4π(sum(areas)/[sum (perimeters)]^2^). In the same analysis, the number of Golgi elements/cell was quantified. Statistical significance was calculated with Kruskal-Wallis test or Mann-Whitney U test with Prism (GraphPad). Experiments were independently repeated at least twice and yielded comparable results. At least 49 cells per treatment condition, per experiment were analyzed. To quantify the FK2 signal, detecting ubiquitinated proteins, at the Golgi, GFP-positive regions (minimum area of 0.05 µm^2^) were identified as above. Integrated density of the FK2 signal within the identified regions was measured in every plane across the z-stack. The sum of the FK2 integrated density was then normalized by the sum of the areas of the GFP-positive regions per cell. Normality of the data was assessed with Shapiro–Wilk, D’Agostino–Pearson and Kolmogorov-Smirnov tests and statistical significance was calculated with Welch’s t-test with Prism (GraphPad). Experiments were independently repeated twice and yielded comparable results. At least 27 cells per treatment condition, per experiment were analyzed. To assess the KLHDC2 signal at the Golgi, a ∼10 µm line was drawn in the juxtanuclear area containing the Golgi, and the intensities of KLHDC2 and the EGFP-fusion proteins along this line were plotted. Measurements were performed on a single plane of a z-stack. Experiments were independently repeated twice and yielded comparable results.

### In situ proximity ligation assay (PLA)

HEK293 Flp-In T-REx cells were plated on coverslips coated with Poly-D-Lysine. After the indicated treatments, cells were fixed using 4% formaldehyde in PBS for 15 min and permeabilized with 0.1 % Triton X-100 in PBS for 10 min at RT. After blocking with Duolink® Blocking Solution for 1 h at 37 °C, cells were incubated in primary antibody solution containing anti-FAF2 (rabbit) and anti-HA tag (mouse) antibodies diluted in Duolink® Antibody Diluent for 1 h at 37 °C. Cells were then washed twice for 5 min with wash buffer A (10 mM Tris-HCl, 150 mM NaCl, 0.05% Tween 20, pH 7.4), followed by incubation for 1 h at 37 °C with secondary antibody solution. Secondary antibodies were oligonucleotide-conjugated PLA probes, Duolink® PLUS (anti-rabbit) and Duolink® MINUS (anti-mouse) PLA probes, diluted in Duolink® Antibody Diluent. Ligation of connector oligonucleotides and the polymerization reaction with fluorescently labeled oligonucleotides were performed with the Duolink® In Situ Detection Reagents Red kit (Sigma-Aldrich, DUO92008) according to manufacturer’s instructions.

### TUBE pull-down

GST-TUBE-His_6_ was expressed in *Escherichia coli* BL21 (DE3) codon plus RIL cells (Novagen) induced with 250 μM isopropyl β-D-1-thiogalactopyranoside for 4 h at 30 °C. Cells were lysed by sonication in 50 mM NaH_2_PO_4_, 300 mM NaCl pH 8.0. The clarified cell lysate was applied to a HisTrap HP chromatography column (Cytiva, GE17-5247). GST-TUBE-His_6_ was eluted in lysis buffer containing 150 mM imidazole. The buffer was exchanged to 20 mM Tris pH 8.0, 100 mM NaCl, 4 mM DTT, 5% glycerol and the concentrated protein was stored at −70 °C until further use. For the TUBE pulldowns, HEK293 Flp-In T-REx GRASP55-HA-EGFP-HT2 and HEK293 Flp-In T-REx TMEM115-HA-EGFP-HT2 (pre-treated with cycloheximide for 1 h) were grown and treated in 10-cm dishes. Cells were washed with cold PBS and lysed in 20 mM Na_2_HPO_4_, 20 mM NaH_2_PO_4_, 1% Triton X-100, and 2 mM EDTA, supplemented with 50 µg/ml purified GST-TUBE-His_6_ protein, cOmplete protease inhibitor cocktail (Roche, 11836145001) and 0.5 mg/ml N-Ethylmaleimide. Cell lysates were clarified by centrifugation at 16,000 g for 20 min at 4 °C, and the supernatant was incubated with Glutathione Sepharose beads (Cytiva, GE17-0756-05) for 2 h at 4 °C. The beads were washed 4x with cold PBS + 0.1% Tween. For pull-downs from HEK293 Flp-In T-REx GRASP55-HA-EGFP-HT2-expressing cells, proteins were eluted by boiling the beads in 1x SDS-PAGE sample buffer (50 mM Tris pH 6.8, 2% SDS, 10% glycerol, 1% β-mercaptoethanol). For pull-downs from HEK293 Flp-In T-REx TMEM115-HA-EGFP-HT2-expressing cells, proteins were eluted by incubating the beads in elution buffer (50 mM Tris, 10 mM L-Glutathione reduced and 0.1% Triton X-100, pH 8.3) for 10 min at RT. Input and elution fractions were mixed with 2x urea sample buffer (50 mM Tris pH 6.8, 2% SDS, 10% glycerol, 4 M urea and 2% β-mercaptoethanol).

### GFP-trap pull-down

For GFP-trap pull-downs, HEK293 Flp-In T-REx GRASP55-HA-EGFP-HT2 and HEK293 Flp-In T-REx TMEM115-HA-EGFP-HT2 were grown and treated in 15-cm dishes. Cells were washed with cold PBS and lysed in 10 mM Tris pH 7.5, 150 mM NaCl, 0.5 mM EDTA, 0.5% Triton X-100 containing cOmplete protease inhibitor cocktail. Cell lysates were clarified by centrifugation at 17,000 g for 20 min at 4°C, and the supernatant was incubated with GFP-trap beads (ChromoTek) for 1 h at 4 °C. The beads were washed 3x with cold wash buffer (10 mM Tris pH 7.5, 150 mM NaCl, 0.5 mM EDTA, 0.05% Triton X-100). For pull-downs from HEK293 Flp-In T-REx cells expressing GRASP55-HA-EGFP-HT2, proteins were eluted by boiling the beads in 2x SDS-PAGE sample buffer. For pulldowns of HEK293 Flp-In T-Rex cells expressing TMEM115-HA-EGFP-HT2, proteins were eluted from the beads by incubating them for 1 min with 200 mM Glycine pH 2.5 followed by pH neutralization with 1 M Tris pH 10.4. Samples were analyzed by western blot (upon mixing with 2x urea buffer) or by mass spectrometry.

### Western blotting

Samples were applied to SDS-PAGE gels and transferred to nitrocellulose membranes. Blocking was performed in 5% non-fat dry milk in TBS-T (TBS containing 0.1% Tween-20). Membranes were incubated with primary antibodies diluted in 3% BSA in TBS-T for 1 h at RT or overnight at 4 °C. HRP-conjugated secondary antibodies were diluted in TBS-T and membranes were incubated in the solution for 45 min at RT. Chemiluminescence signals were visualized with ECL Start and ECL Prime detection reagents (Cytiva, RPN3243 and RPN2232) at a ChemiDoc MP Imaging System (BioRad).

### Sample preparation for Mass Spectrometry

#### Sample preparation for enriched proteome samples

Eluted protein samples from the GFP trap pull-down were taken up in SP3 Lysis buffer (final concentrations; 1% SDS, 10 mM TCEP, 40 mM chloroacetamide, 50 mM HEPES pH 8.0), heated up to 90 °C for 5 min and cleared by centrifugation. The subsequent sample preparation for proteomic analysis was performed using the single-pot, solid-phase-enhanced sample-preparation (SP3) strategy^98^. Supernatant was transferred to fresh tube (∼50 µL volume). Then, 3 µL of a 50 µg/µL 1:1 mixture of hydrophilic (#45152105050250) and hydrophobic (#65152105050250) carboxylate modified Sera-Mag SpeedBeads (Cytiva), that were washed twice with MS-grade water, were added to the samples. The next steps were carried out at room temperature unless noted otherwise. Afterwards, the samples were mixed shortly (1 min, 1000 rpm) and collected by short centrifugation (10 s, 200 g). Next, protein binding was induced by the addition of an equal volume of pure ethanol (10 min, 1000 rpm). After a short centrifugation step (10 s, 200 g) beads were collected by placing the reaction vessel on a magnetic stand. Beads were allowed to migrate towards the magnet for at least 5 min before the supernatant was removed. The beads were then taken up in 180 µL 80% (v/v) ethanol and transferred to a fresh multiwell plate. Subsequently the beads were washed four times with 180 µL 80% (v/v) ethanol prior to the addition of 100 µL digestion enzyme mix (0.6 μg of trypsin (V5111; Promega) and 0.6 µg LysC (125-05061; FUJIFILM Wako Pure Chemical) in 25 mM ammonium bicarbonate). Samples were incubated at 37 °C for 19 h while shaking (1300 rpm). On the next day, the samples were briefly centrifuged (10 s, 200 g) and placed on a magnet for 5 min. The clear solution containing the tryptic peptides was transferred to a fresh multiwell plate. The beads were taken up in 47 µL 25 mM ammonium bicarbonate and incubated while shaking (10 min, 1000 rpm). The plate was then placed on a magnetic stand and after 5 min the cleared supernatant was collected and combined with the recovered first peptide mix, followed by the addition of formic acid (FA) to a final concentration of 2% (v/v) for trypsin inactivation.

#### Sample clean-up for LC-MS/MS

All digests were desalted on home-made C18 StageTips^99^ containing two layers of an octadecyl silica membrane (CDS Analytical). All centrifugation steps were carried out at room temperature. The StageTips were first activated and equilibrated by passing 50 μL of methanol (600 g, 2 min), 80% (v/v) acetonitrile (ACN) with 0.5% (v/v) FA (600 g, 2 min) and 0.5% (v/v) FA (600 g, 2 min) over the tips. Next, the acidified tryptic digests were passed over the tips (800 g, 3 min). The immobilized peptides were then washed with 50 μL and 25 μL 0.5% (v/v) FA (800 g, 3 min). Bound peptides were eluted from the StageTips by application of two rounds of 25 μL 80% (v/v) ACN with 0.5% (v/v) FA (800 g, 2 min). Peptide samples were then dried using a vacuum concentrator (Eppendorf) and the peptides were dissolved in 15 μl 0.1% (v/v) FA prior to analysis by MS.

### LC-MS/MS settings

#### LC-MS/MS Analysis

LC-MS/MS analysis of peptide samples was performed on an Orbitrap Fusion Lumos mass spectrometer (Thermo Scientific) coupled to a Vanquish Neo ultra high-performance liquid chromatography (UHPLC) system (Thermo Scientific) that was operated in the one-column mode. The analytical column was a fused silica capillary (inner diameter: 75 μm, outer diameter: 360 µm, length: 28 cm; CoAnn Technologies) with an integrated sintered frit packed in-house with Kinetex 1.7 μm XB-C18 core shell material (Phenomenex). The analytical column was encased by a PRSO-V2 column oven (Sonation) and attached to a nanospray flex ion source (Thermo Scientific). The column oven temperature was set to 50 °C during sample loading and data acquisition. The LC was equipped with two mobile phases: solvent A (2% ACN and 0.2% FA, in water) and solvent B (80% ACN and 0.2% FA, in water). All solvents were of UHPLC grade (Honeywell). Peptides were directly loaded onto the analytical column with a maximum flow rate that would not exceed the set pressure limit of 950 bar (usually around 0.5 – 0.6 μL/min) and separated on the analytical column by running a 105 min gradient of solvent A and solvent B at a flow rate of 300 nL/min (start with 3% (v/v) B, gradient 3% to 6% (v/v) B for 5 min, gradient 6% to 29% (v/v) B for 70 min, gradient 29% to 42% (v/v) B for 15 min, gradient 42% to 100% (v/v) B for 5 min and 100% (v/v) B for 10 min). The mass spectrometer was controlled by the Orbitrap Fusion Lumos Tune Application (version 4.1.4244) and operated using the Xcalibur software (version 4.7.69.37). **Table S3** gives an overview of the most important MS settings:

RAW spectra were submitted to a closed MSFragger (version 4.1.)^100^ search in Fragpipe (version 22)^101^ using the “LFQ-MBR” workflow (label-free quantification and match-between-runs; default settings were used unless otherwise stated). RAW files were listed in the “Input LC-MS Files” section and experiment set “by file name”. As “Data Type” we kept the default “DDA” (data dependent acquisition). The MS/MS spectra were searched against a custom database generated in Fragpipe (1. 2025-07-24-decoys-contam-UP000005640_9606_OPPG_plusSOI.fasta.fas (41310 entries)) containing the Uniprot *Homo sapiens* reference proteome (UP000005640_9606_OPPG.fasta; 20067 entries), the sequence of interest (TMEM115–HA-tag–EGFP-HaloTag2), contaminants and decoys (the last two appended and generated by Fragpipe in the Database section). MSFragger searches allowed oxidation of methionine residues (16 Da; 3) and acetylation of the protein N-terminus (42 Da;1) as variable modification (first value in brackets refers to the molecular weight of the modification, second value to the maximum number of occurrences per peptide). Carbamidomethylation on Cysteine (57) was selected as static modification. Enzyme specificity was set to “Trypsin specific”. The initial precursor and fragment mass tolerance was kept at ±20 ppm. Mass calibration and parameter optimization was selected. Validation of peptide spectrum matches was done using MSBooster^102^ using DIA-NN^103^ for RT and spectra prediction. Peptide spectra matches (PSM) were validated using percolator with a minimum probability setting of 0.5. Protein inference was performed using ProteinProphet (part of Philosopher version 5.1.1). The final reported protein FDR was 0.01 (based on target-decoy approach). Protein quantification was performed with IonQuant (version 1.10.27). Add MaxLFQ (min ions 1), MBR (FDR 0.01) and normalization of intensities across runs was selected. Unique and razor peptides were allowed. Advanced options were kept at default. Further analysis and filtering of the results was done in Perseus v1.6.10.0^104^. For quantification, related biological replicates were combined into categorical groups, and only proteins with at least 4 valid values in at least one categorical group were analyzed. Comparison of protein group quantities (relative quantification) between different MS runs is based solely on the LFQs as calculated by IonQuant (MaxLFQ algorithm). Data were visualized using Instant Clue^96^.

