## Supplementary Figures for "Selective proteasomal degradation from the Golgi apparatus membrane"

**Content**

**Supplementary Figures 1-5, Supplementary Table 3**

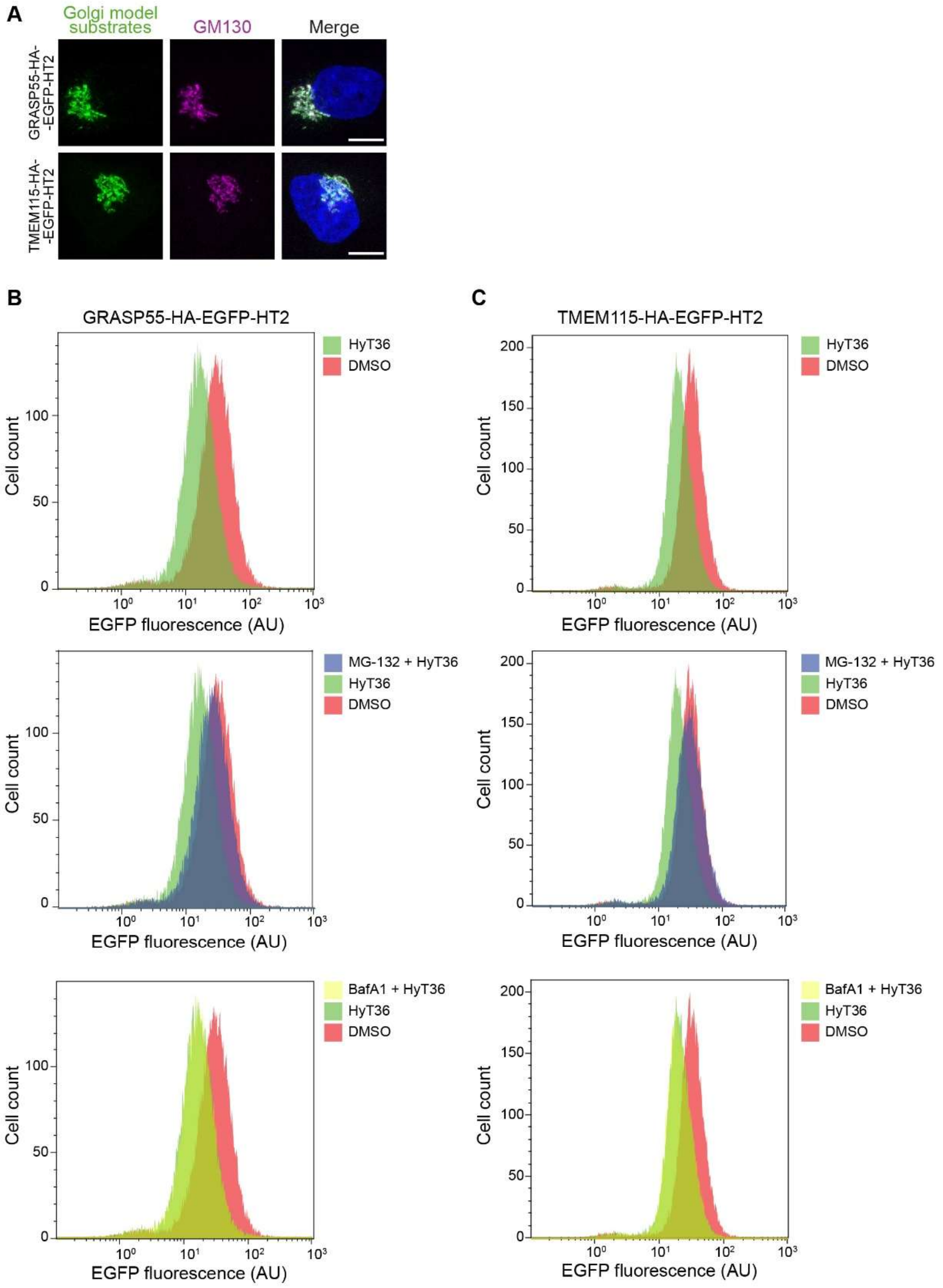
A
Golgi model substrates
GM130
Merge
GRASP55-HA-EGFP-HT2
TMEM115-HA-EGFP-HT2
B
GRASP55-HA-EGFP-HT2
C
TMEM115-HA-EGFP-HT2
HyT36
DMSO
MG-132 + HyT36
BafA1 + HyT36
Cell count
EGFP fluorescence (AU)

**Figure S1. Characterization of cell lines expressing Golgi model substrates.** (**A**) Representative maximum intensity projections of confocal stacks showing HeLa Flp-In T-REx cells expressing GRASP55-HA-EGFP-HT2 or TMEM115-HA-EGFP-HT2. (**B**) Representative flow cytometry raw data showing EGFP fluorescence in arbitrary units (AU) and according cell counts for HEK293 cells expressing GRASP55-HA-EGFP-HT2 exposed to the indicated treatments for 5 h following a 1 h cycloheximide pre-treatment. (**C**) Same analysis as in (B) performed with HEK293 cells expressing TMEM115-HA-EGFP-HT2.

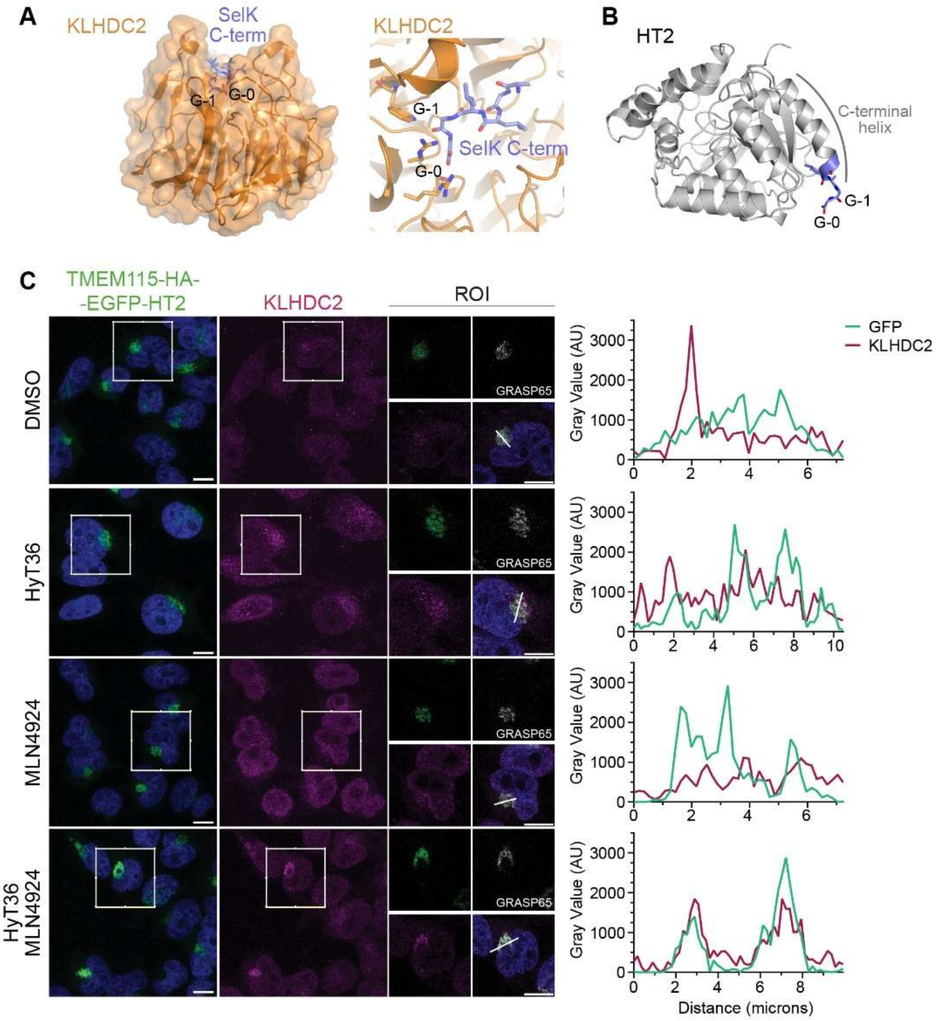


**Figure S2. The CRL2$^{KLHDC2}$ ligase targets HT2 fusion proteins.** (**A**) Crystal structure of KLHDC2 in complex with the six most C-terminal residues of its substrate protein SelK (PDB ID: 6DO5 chain A and C). Surface representation shows binding of the substrate deep in the KLHDC2 propeller. Zoom-in shows the extended conformation of the SelK peptide. (**B**) Alphafold3 model of the HaloTag2 sequence used in this study ending in -GLAGG. These last five residues are highlighted in purple. (**C**) *Left panel:* Overview images show representative maximum projections of confocal image stacks. Representative confocal sections of HeLa Flp-In T-REx cells expressing TMEM115-HA-EGFP-HT2. Following the indicated treatments for 5 h cells were immunostained for KLHDC2 and the Golgi marker GRASP65. *Right panel:* Quantification of gray value along the line shown in the region of interest (ROI). Scale bar 10 µm. Arbitrary units (AU).

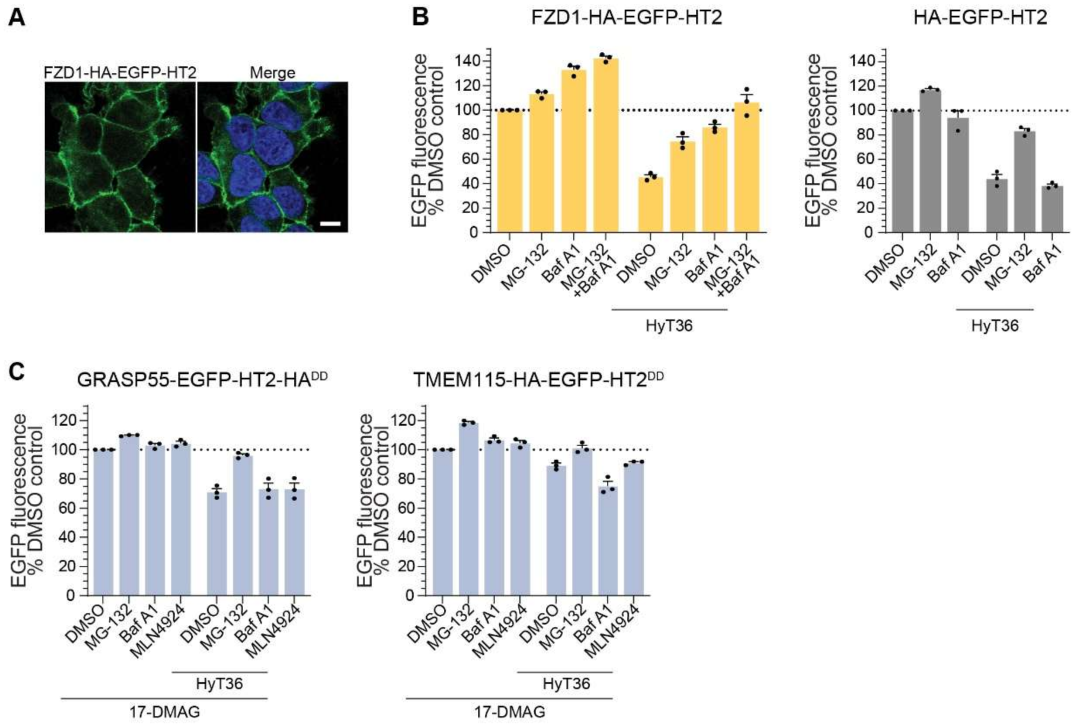


**Figure S3. Degradation profile of EGFP-HT2 fusion proteins.** (**A**) Representative maximum intensity projections of confocal stacks showing HEK293 cells expressing FZD1-HA-EGFP-HT2. Nuclei (blue) were stained with Hoechst. Scale bar, 10 µm. (**B**) Levels of FZD1-HA-EGFP-HT2 and HA-EGFP-HT2 as determined by flow cytometry. Indicated treatments were applied for 5 h following a 1 h pre-treatment with cycloheximide. Data represent three independent experiments. Graphs show mean + S.D. (**C**) Levels of Golgi model substrate constructs ending in -DD as determined by flow cytometry. All conditions include the Hsp90 inhibitor 17-DMAG. Indicated treatments were applied for 8 h following a 1 h pre-treatment with cycloheximide. Data represent three independent experiments. Graphs show mean + S.D.

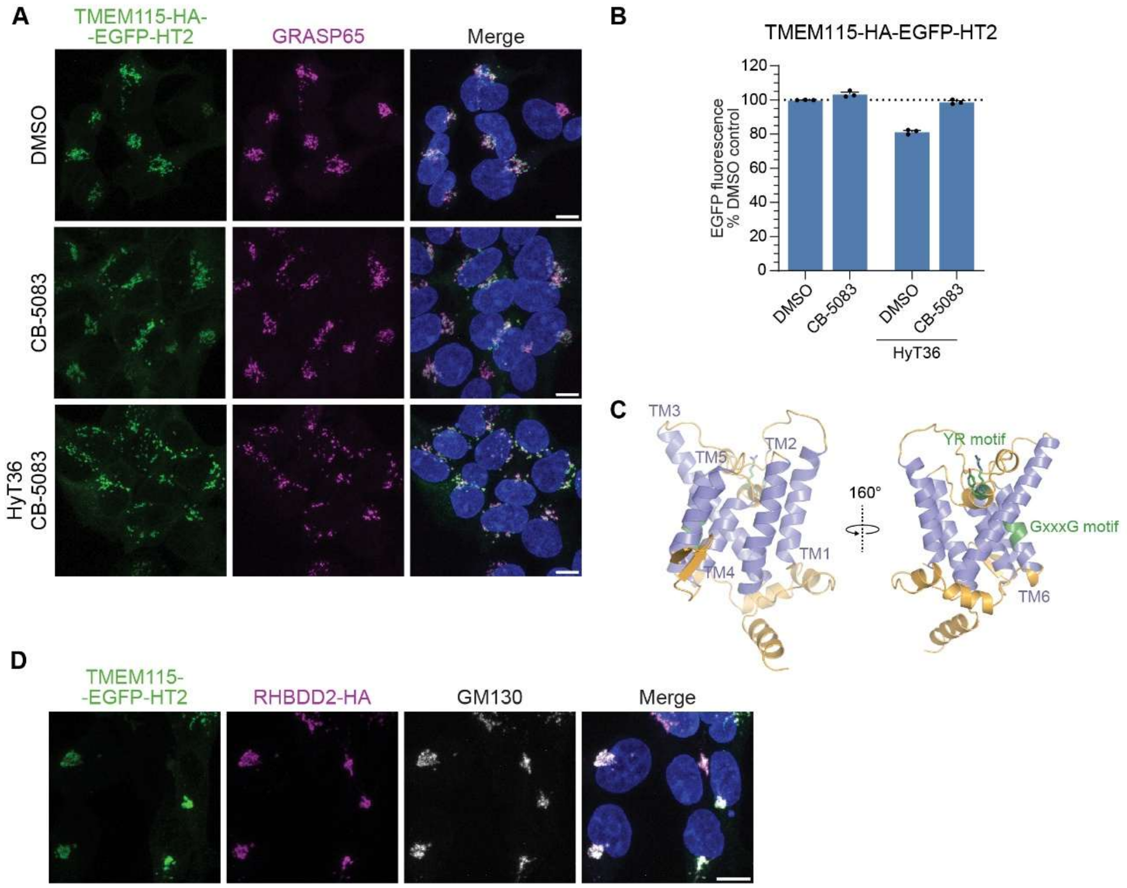


**Figure S4. Localization of Golgi model substrates and RHBDD2.** (**A**) Representative maximum intensity projections of confocal stacks showing the localization of TMEM115-HA-EGFP-HT2 in HEK293 cells treated with the p97 inhibitor CB-5083 and/or HyT36 for 5 h. Immunostaining of GRASP65 illustrates the Golgi. (**B**) Levels of TMEM115-HA-EGFP-HT2 in HEK293 cells determined by flow cytometry under treatment conditions reflecting the pull-down set-up in Figure 5. Indicated treatments were applied for 3 h following a 3 h cycloheximide pre-treatment. Graph shows mean + S.D. (**C**) Alphafold model of RHBDD2 (AF-Q6NTF9-F1-model_v6). Flexible N-terminal and C-terminal residues (1-16 and 250-364) omitted for clarity. Transmembrane helices (magenta) and conserved motifs (YR, GxxG) are highlighted. (**D**) Representative maximum intensity projections of confocal stacks showing the localization of TMEM115-EGFP-HT2 upon doxycycline induction and RHBDD2, constitutively expressed, in HEK293 cells. Immunostaining of GM130 illustrates the Golgi.

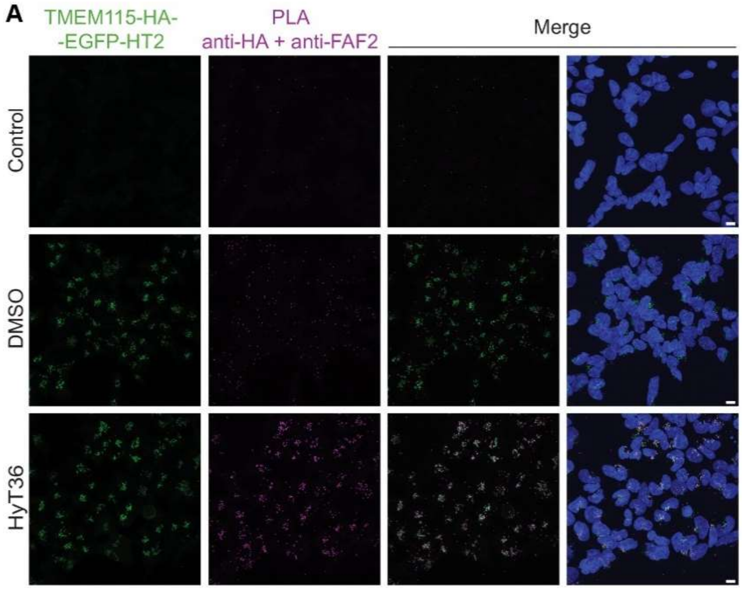


**Figure S5. Overview of PLA imaging data related to Figure 6B.** Proximity ligation assay (PLA) using FAF2- and HA-antibodies in three conditions. In control conditions, the expression of TMEM115-HA-EGFP-HT2 integrated into HEK293 cells was not induced. Doxycycline-induced HEK293 cells were treated with DMSO or HyT36 for 3 h following a 3 h cycloheximide pre-treatment.

### Table S1. MAGeCK-RRA analysis of sgRNAs and genes included in the CRISPR/Cas9 screen, comparing top25 and bottom25

### Table S2. Proteomics raw data of GFP trap pull-down of TMEM115-HA-EGFP-HT2 with CB-5083 vs CB-5083 + HyT36

### Table S3. MS Settings

| Project | MS | general | MS1 | MS2 | MS2 | MS3 | Comments; special settings |
|---|---|---|---|---|---|---|---|
| ACE_1003 | Lumos | Tune v4.1.4244<br>Xcalibur v4.7.69.37<br>SII: 1.7.0.468<br>Gradient: 105 min | Analyzer: FT<br>Res.: 240000<br>SR: 375 - 1500<br>AGC: Standard<br>AGC abs.: 400000<br>AcT: 50 ms<br>RF: 30<br>SF: --<br>DDM: CT/3sec | Analyzer: IT<br>Res./ScR: -/rapid<br>SR: Auto<br>AGC: 300%<br>AGC abs.: 30000<br>AcT: auto<br>CS: +2 to +7<br>IsM: Q<br>IsW: 1.6<br>Frag.: sHCD<br>NCE: 20, 30, 45 | | | classic orbitrap experiment: MS1 in Orbitrap at high resolution and data dependent MS2 in Iontrap rapid scan rate. Dynamic exclusion enabled (exclude after n times=1; Exclusion duration (s)= 20; mass tolerance= ± 10ppm)<br><br>Intensity Threshold: 5000<br>Ion transfer Tube Temp: 230 °C<br>Ion Source Voltage: 2500 V |

**FT**= Fourier Transform (Orbitrap); **IT**= Iontrap; **Q**= Quadrupol; **Res.**= max. Resolution at 200 m/z (Lumos) or 400 m/z (Elite) [FWHM (full width at half maximum)]; **ScR**= scan rate for measurements in the IT; **SR**= scan range [m/z]; **AGC**= automatic gain control, max number of acquired ions per measurement; **AcT**= max. Ion acquisition time [ms]; **CS**= charge states used for fragmentation; **IsM**= Isolation mode (Q or IT), MS2 isolation and further is only done in IT; **IsW**= Isolation window [m/z], value followed by scan mode the isolation is based on (MS1, MS2 …) **Frag.**= Fragmentation method; **HCD**= Higher-energy collisional dissociation; **CID**= Collision-induced dissociation; **ETD**= Electron-transfer dissociation**; EThcD=** Electron-Transfer/Higher-Energy Collision Dissociation; **sHCD**= stepped HCD**; NCE**= normalized collision energy; **cycles**: number of MSn recorded or max cycle time; **RF**= RF Lens [%]; **SF**= Source Fragmentation [V]; **DDM**: Data dependent Mode (cycle time in seconds, CT/[s] or number of scans, NS); **NS**= Number of data dependent scans
